# Transitions in paternal dominance regulate offspring growth and metabolic transcription

**DOI:** 10.1101/443317

**Authors:** Joseph W. Cauceglia, Adam C. Nelson, Nimrod D. Rubinstein, Shweta Kukreja, Lynsey N Sasso, John A. Beaufort, Oliver J. Rando, Wayne K Potts

## Abstract

Parental effects are an important source of adaptive traits. By contrast, parental effects failing to regulate offspring phenotype to fit current conditions could be deleterious. Although adaptive parental responses to single cues have been identified, we lack an understanding of the reversibility of parental effects across breeding cycles in a fluctuating environment. Social status of parents can occasionally fluctuate and, in turn, influence high-fitness pathways available to offspring. We show that social competition status results in robust parental effects on growth in mice. Dominant males produce faster growing offspring because of status related cues, not genetic associations. The timing, effect-size, and sex-specificity of this paternal effect are modulated by maternal experience. We experimentally demonstrate that status-ascending males produce heavier sons than before, and status-descending males produce lighter sons than before. Paternal status predicts genome-wide transcription in the liver, including transcriptional networks controlling xenobiotic and fatty acid metabolism, and oxidative phosphorylation. Our study demonstrates that paternal social status reversibly conditions offspring growth in naturalistic environments.

## INTRODUCTION

Parental effects are an important source of phenotypic variation. Parental effects can be adaptive in the sense that they better prepare offspring for current environmental challenges (Badyaev & Uller, 2009). For placental mammals, pre-and post-natal development is highly dependent on the mother, and many adaptive parental effects have been found to have a maternal origin (Mousseau *et al.*, 2009). Although fathers contribute less to development, they too can influence offspring traits in significant ways (Rando, 2012). Evolutionary theory predicts conflicts over maternal and paternal regulation of offspring development (Trivers, 1974). While females must balance the current benefits of reproduction with long-term survival and reproduction, males can benefit from inducing females to disproportionately increase investment in offspring (Stearns, 1992). Determining how maternal and paternal effects are balanced during development is an important question in biology.

In a fluctuating environment, adaptive parental effects should be reversible from one reproductive episode to the next, depending on the time scale of environmental variation; failure to appropriately program offspring phenotype with current conditions could produce a deleterious outcome (DeWitt *et al.*, 1998). Many phenotypically plastic traits fall somewhere between being nearly irreversible or instantaneously reversible, depending on factors such as response time, pattern of exposure, and the quality of information available (Gabriel *et al.*, 2005). Accordingly, some mammalian parental effects “carry over” for one to several generations after the stimulus has been removed (Cropley *et al.*, 2012; Dias & Ressler, 2014; Bošković & Rando, 2018). If conditions reliably change on a time-scale that is longer than the breeding cycle, but shorter than reproductive lifetime, then evolution should favor fast and reversible effects. One prediction, then, is that parents should be able to produce offspring with multiple phenotypes across breeding cycles (Lacey, 1998). Although adaptive maternal and paternal responses to single cues have been identified (Bonduriansky & Head, 2007; Ducatez *et al.*, 2012; Stein & Bell, 2014; Vallaster *et al.*, 2017), their reversibility under fluctuating conditions is understudied (Jensen *et al.*, 2014), particularly in mammals.

For many animals, the social environment is organized around dominance hierarchies. Individual position within the hierarchy is a major determinant of fitness as well as physiology (Wilson, 1975; Creel *et al.*, 2012). Though generally stable, hierarchies undergo periods of instability as individuals gain and lose competitive ability (Sapolsky, 2005; Shizuka & McDonald, 2015). In house mice, social groups are organized around the territories of dominant males, who continuously have to defend their position against competitors (Bronson, 1979). Although less understood, female mice also compete for territories and reproduction in cooperative groups (Harrison *et al.*, 2018). The territories of dominant males are rich in resources and protected against predators and competitors, while nondominant males live in vulnerable, low-quality territories with other nondominant males. Thus, offspring of dominant fathers face radically different environmental conditions than those born to nondominants; in turn, sons and daughters employ territory-dependent developmental strategies as they mature (Gerlach, 1990; 1996).

Here, we set out to systematically dissect the role of maternal and paternal effects on trait variability in a fluctuating social environment. We previously established that parental effects could contribute to rapid adaptation to the social environment (Nelson *et al.*, 2013a; Nelson *et al.*, 2013b). This finding prompted us to use reciprocal crosses to quantify maternal and paternal effects of social competition. Here we show that that offspring body weight is acutely susceptible to parental effects that operate at different developmental timepoints. Intriguingly, paternal effects can be a stronger predictor of growth rate than maternal effects, and are reversible on the order of weeks. Finally, paternal social status is associated with gene-expression in offspring, including pathways that control metabolism of glucose, amino acids, lipids, and iron.

## RESULTS

### Parental social competition and dominance hierarchy

To investigate parental effects of social experience, we employed a breeding strategy where adult mice experience social competition in seminatural enclosures prior to breeding under controlled, monogamous conditions (Methods). In seminatural enclosures, males and females compete for high-quality territories providing a defendable shelter and food; in contrast, low-quality territories are exposed and associated with communal feeding sites (Nelson *et al.*, 2013a; Nelson *et al.*, 2013b). Socially dominant males are characterized as having a near-exclusive occupancy of high-quality territories; in some cases, a single dominant male will control multiple territories (Fig. 1B, C). Dominant males also produce more offspring (Nelson *et al.*, 2013b). Although female mice form distinct, spatially organized social interactions, there is not a clear dominance hierarchy among them (Fig. 1B, C).

**Figure 1.**
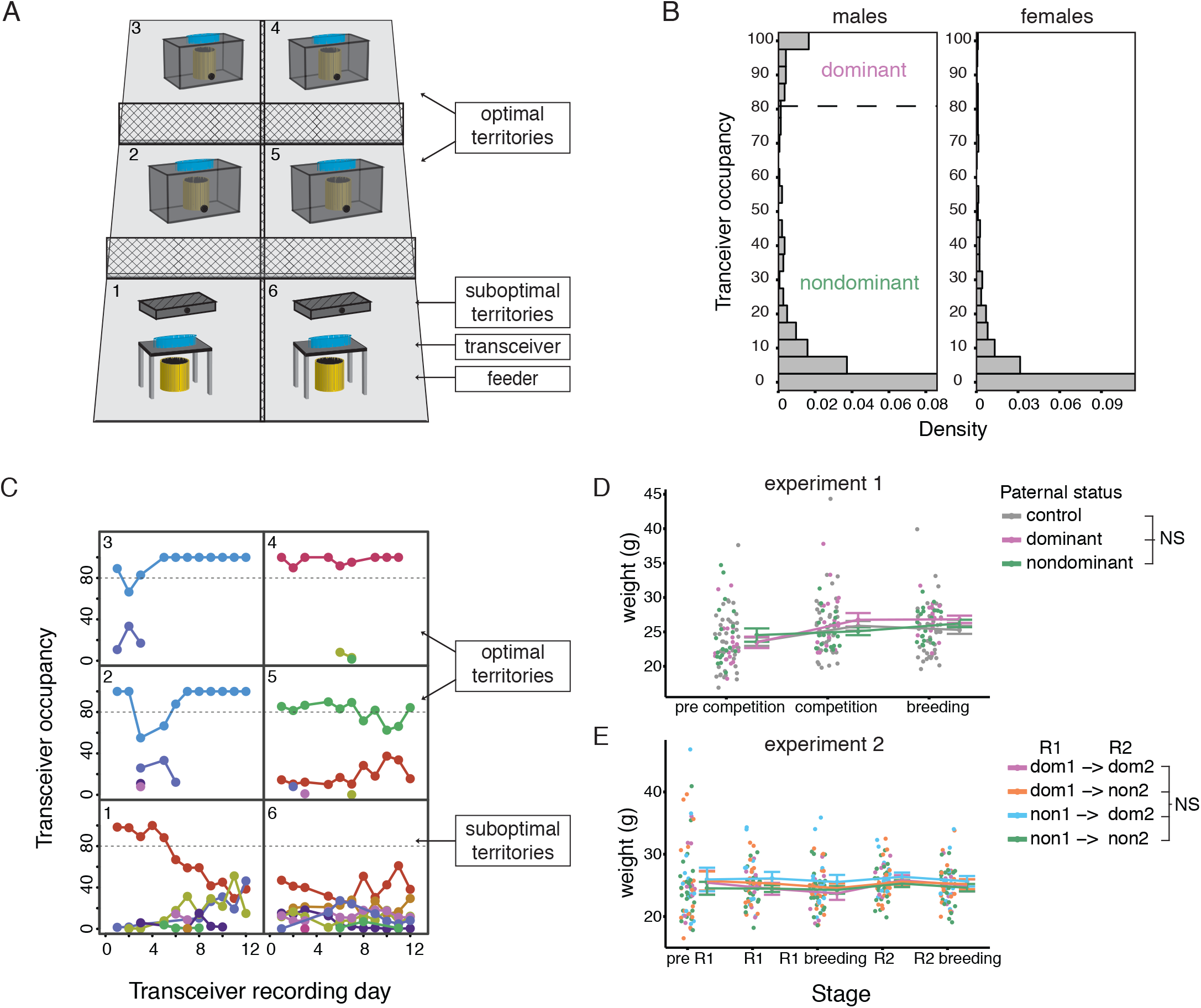
Identification of social dominance hierarchies. (A) Schematic of seminatural enclosures. RFID transceivers are placed on the food-source at suboptimal (sections one & six) and optimal (sections two to five) territories, and tagged mice are monitored for their occupancy of the territories. (B) Density distributions of transceiver occupancy. Left: dominant males are defined by 80% occupancy of a single territory. Right: females do not display social dominance in the same way as males. N = 93 males, N = 123 females. (C) RFID reads of male mice in one enclosure; each color represents a single male. Dominant males were detected in territories two and three (blue line), territory four (red line), and territory five (green line). The remaining males observed in territories 1 & 6 were nondominant. (D) Body weight is not associated with social dominance in the paternal males used in the two breeding experiments described hereafter. Experiment 1 (top): mean weight (and SEM) of control (N = 42), dominant (N = 25), and nondominant (N = 21) sires before and after hierarchy establishment, and during breeding. Experiment 2 (bottom): mean weight (and SEM) of sires that maintained status (dom->dom (N = 19) & non->non (N = 25) or reversed status (dom->non (N = 19) & non->dom (N = 15)) over two rounds of competition. Experiment 2 stages: before first round of competition (R1); during first round of competition; during first round of controlled breeding; during second round of competition (R2); during second round of controlled breeding.

We addressed the role of parental experience of social competition and hierarchy on offspring growth in two experiments. In experiment 1, we determined the relative effects of maternal competition and paternal social dominance status compared to non-competition controls. In experiment 2, we determined paternal effects of social dominance status transitions. Although we report below parental effects on offspring weight, we note that for animals from the social competition cohort (i.e., the parental generation) social status was unassociated with body weight in males (Fig. 1D) and females (linear mixed models, P > 0.05). These results indicate that when introduced to seminatural enclosures, mice compete for territorial resources, males form a clear division between dominant and nondominant status, and social status is unassociated with body weight.

## EXPERIMENT 1

### Maternal and paternal social experience affects offspring growth throughout development

To identify the relative contribution of maternal and paternal effects of social competition, adult mice (N = 222; 80 males, 142 females) carrying the wild-derived “CNGWLD” genetic background were introduced to seminatural enclosures (N = 8) (Methods). In parallel, a control, noncompetition group of age-matched mice (N = 104) were assigned to monogamous breeding cages (N = 52 pairs). After an 11-week social exposure, animals were categorized by their condition (competition or control), with competition males further classified as dominant or nondominant (Fig. 2A). A reciprocal F1 cross was then employed to systematically assess parental effects of social experience and status on offspring body weight. All combinations of female and male conditions were bred as monogamous pairs in standard cages for 10 days, after which the male was removed (Fig. 2A). These pairs yielded 56 litters and 420 offspring. Growth rate of the resulting offspring was recorded for 32 weeks. We used linear mixed models (LMM) and best-fit model selection (Akaike information criterion, AIC) to assess parental effects on body weight while accounting fixed and random effects (Methods).

**Figure 2.**
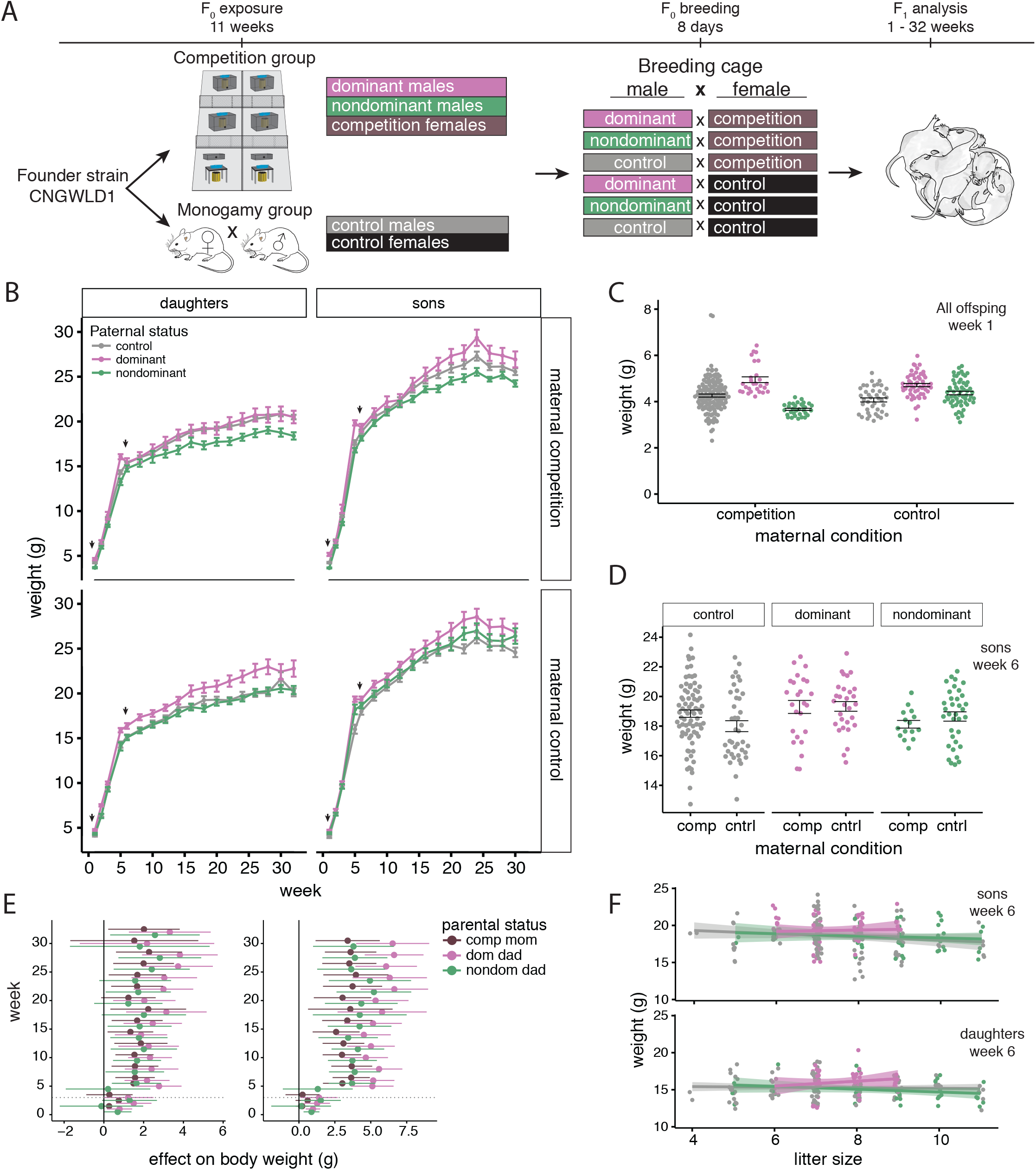
Maternal and paternal effects of social status on offspring growth. (A) Design of Experiment 1. Parental animals from the CNGWLD strain experienced either 11 weeks of competition in seminatural enclosures (dominant males, nondominant males, and competition females) or monogamous mating (control males and females). Males and females were mated monogamously in cages in a full factorial F1 cross (N = 80 pairs; 56 litters). Offspring weight was analyzed for 32 weeks. (B) Maternal and paternal effects of social competition on the weight of daughters (N = 198) and sons (N = 222). Body mass trajectories (mean and SEM) of daughters (left; to 32 weeks of age) and sons (right; to 30 weeks of age). Maternal effects of social competition (top row) and noncompetition control (bottom row). (B) Black arrows indicate timepoints expanded in panel C and D. (C) Relationship between maternal (competition or control) and paternal (control, dominant or nondominant) status and offspring weight at postnatal week one in all offspring (top). Each point represents a single mouse with group means and SEM (N = 343 offspring). (D) Relationship between paternal status (control, dominant or nondominant) for each maternal condition (competition or control) and offspring weight at week six in sons. Each point represents a single mouse with group means and SEM (N = 222 sons). (E) Forest plots show the association between parental social status and body mass of daughters (left) and sons (right) while controlling for other factors; data are coefficients from LMM at each time point separately. Means and 95% confidence intervals indicate the effect size on body weight due to maternal social competition and paternal social dominance status relative to monogamous control parents (zero line). Dotted lines at week 3 indicate time of weaning. (F) Paternal effects on the relationship between litter size and mean, per-litter weight of sons (top) and daughters (bottom) at week six. Shown are best fit lines and SEM.

Full models were the best-fit at both developmental stages (Table 1), indicating that paternal condition, maternal condition, age, litter size, litter sex ratio, and their interactions are primary determinants of offspring growth. Generally, parental effects of social competition on sons and daughters were robust throughout development (Fig. 2, Tables 1-2). Overall, the strongest growth-promoting parental effect was having a dominant father (Fig. 2B-E, Table 2). In addition, relative to noncompetition mothers, competition mothers produced heavier offspring (Fig. 2D & E). The effect of paternal nondominance was dependent on maternal competition; nondominant fathers produced lighter offspring when mated with competition mothers but not control mothers (Fig. 2C & D, Table 2). Details of these effects are described below.

**Table 1.** AIC selection of best fitting models for Experiment 1 and Experiment 2.

|  | Pre-puberty |  | Post-puberty |  |
| --- | --- | --- | --- | --- |
| | AIC | $\Delta$ AIC | AIC | $\Delta$ AIC |
| **Experiment 1** |  |  |  |  |
| Full Model: sex, litter size, litter proportion male, father's status, mother's status | 5609.32 | 0.00 | 22066.64 | 0.00 |
| Sex removed | 6344.82 | +735.49 | 22804.67 | +738.03 |
| Litter size removed | 5668.67 | +59.34 | 22223.78 | +157.13 |
| Proportion male removed | 5696.85 | +87.52 | 22149.07 | +82.43 |
| Father's status removed | 5703.41 | +94.08 | 22247.09 | +180.45 |
| Mother's condition removed | 5684.33 | +75.00 | 22154.24 | +87.6 |
| Direct effects only (no interactions) | 6228.11 | +618.78 | 22570.64 | +503.99 |
| **Experiment 2** |  |  |  |  |
| Full Model: sex, litter size, litter proportion male, father's round 1 status | 6280.27 | 0.00 |  |  |
| Sex removed | 7062.74 | +782.47 |  |  |
| Litter size removed | 6300.38 | +20.11 |  |  |
| Proportion male removed | 6303.16 | +22.89 |  |  |
| Father's status removed | 6286.45 | +6.17 |  |  |
| Direct effects only (no interactions) | 6935.03 | +654.76 |  |  |

Selected examples of AIC comparisons. In all cases the full model had better AIC score than any models with one or multiple terms removed. Abbreviations: AIC = Akaike information criterion. Random effects structures were similarly determined with stepwise AIC selection, and required control for repeated measures (offspring ID) and sibling relatedness (birth cage).

**Table 2.**
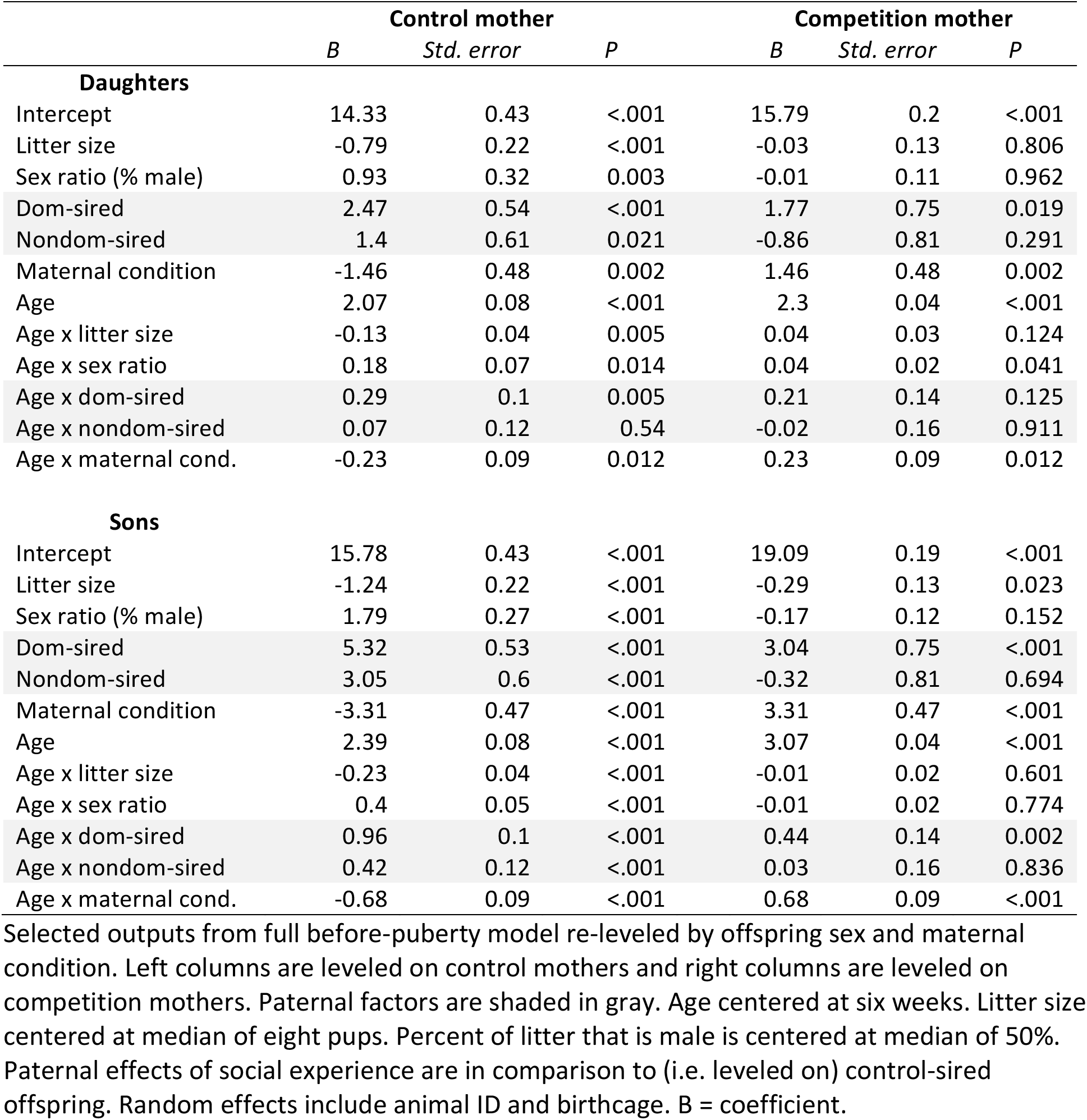
Pre-puberty predictors of offspring weight in Experiment 1. Refers to Figure 2.

### Dominant males produce heavier offspring when paired with control females

We first addressed parental effects of social competition by examining the offspring of control mothers. At the first recorded weight (week one), sons and daughters weighed equivalently and were analyzed together; offspring of dominant fathers were 23% heavier than those of control fathers (Fig 2C; LMM week 1: 0.89 ± 0.29 above 3.87 ± 0.24 g, P = 0.002). At six weeks, dominant-sired daughters were 17% heavier than those of control-sires (Table 2), and dominant-sired sons were 34% heavier than those of control-sires (Fig. 2D, Table 2). Additionally, nondominant-sired sons at six weeks were 19% heavier than those of control-sires (Fig. 2B, Table 2). Thus, control mothers mated with dominant males (and to a lesser extent, nondominant males) produce heavier offspring, especially sons.

We then examined offspring growth rate of control mothers (Tables 2 & SI Table 1). Before puberty, dominant-sired daughters grew 14% faster than control sired daughters (Age x dominant-sire interaction, Table 2 Daughters), while nondominant-and control-sired daughters were equivalent. Dominant-sired sons grew 40% faster, and nondominant-sired sons grew 18% faster than control-sired sons (Age x paternal status interactions, Table 2 Sons). After puberty, the effect of having a dominant father persisted: dominant-sired daughters maintained a 26% faster growth rate than control-sired daughters, and dominant-sired sons grew 27% faster than control-sired sons (Age x paternal status interactions, SI Table 1). In contrast, nondominant-sired sons were no longer heavier than control-sired sons after puberty. These results suggest that paternal effects of social status regulate offspring weight gain throughout development, with dominant fathers increasing the growth of sons being the strongest effect.

### Maternal effects modulate paternal effects on offspring growth

We next examined maternal effects and the relationship between maternal and paternal effects. First, we found that competition mothers produced heavier offspring than control mothers throughout development, though the effect was not as strong as having a dominant father (Fig. 2E). We then evaluated whether the paternal effects observed with control mothers were also observed with competition mothers. Like control mothers, competition mothers had heavier daughters and sons when mated to a dominant male (Fig. 2C, D, E, Table 2). However, competition and control mothers responded differently to nondominant fathers: competition mothers mated to nondominant males produced offspring that were 26% lighter at one week of age than those of control mothers (Fig. 2C; LMM week 1: −1.23 ± 0.45 above 4.66 ± 0.21 g, P = 0.006). Competition mothers also responded differentially to dominant vs. nondominant fathers: dominant-sired offspring were significantly (36%) heavier at one week of age than nondominant-sired offspring (Fig. 2C; LMM week 1; 1.25 ± 0.56 above 3.44 ± 0.40 g, P = 0.026).

We then evaluated maternal effects on the growth rate before puberty (Fig. 2B and Table 2). Competition and control mothers responded differently to control fathers; daughters of competition mothers grew 14% faster (age x maternal condition interaction, SI Table 2 Control-sired), and sons grew 28% faster (age x maternal condition interaction, SI Table 2 Control-sired) than daughters and sons of control mothers. Next, analysis of growth after puberty showed that competition and control mothers responded differently to nondominant fathers: nondominant-sired sons of competition mothers grew 34% slower than those of control mothers (age x nondominant-sired interaction, SI Table 1 Sons). Thus, compared to control mothers, competition mothers invest more in offspring growth, respond favorably to dominant fathers and unfavorably to nondominant fathers, and these responses play out at different developmental timepoints.

Both maternal and paternal competition induced growth-prompting effects in offspring (Fig. 2E, Tables 2 & SI Tables 1-3), prompting us to investigate their relative strength and developmental timing. Effect size analysis of per-week weight gain showed that paternal dominance was the strongest determinant: dominant-sired daughters gained more weight at weeks 2 and 22, and dominant-sired sons gained more weight at weeks 1, 5, and 22 (Supplementary Fig. 1A).

**Table 3.**
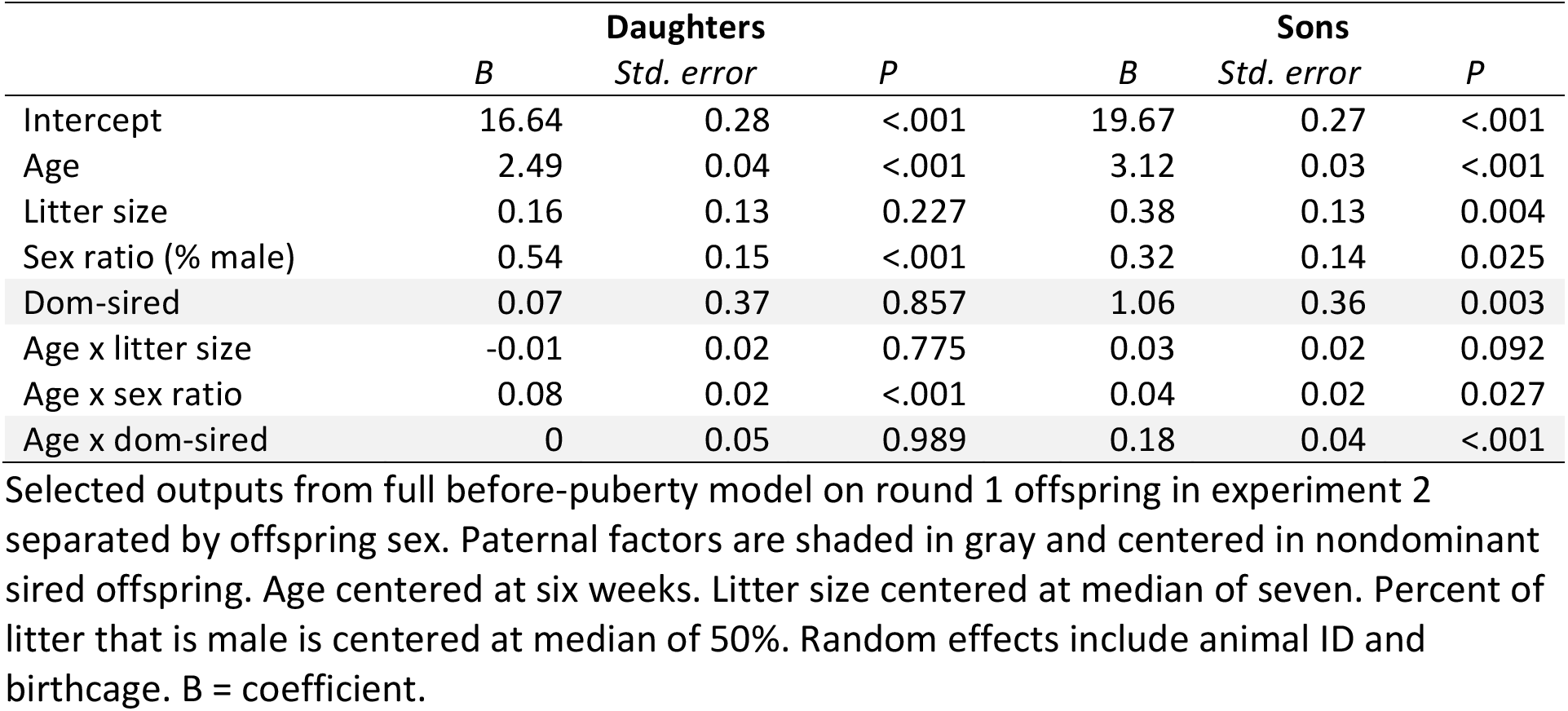
Pre-puberty predictors of offspring weight in Experiment 2. Refers to Figure 3.

|  | Daughters |  |  | Sons |  |  |
| --- | --- | --- | --- | --- | --- | --- |
|  | *B* | *Std. error* | *P* | *B* | *Std. error* | *P* |
| Intercept | 16.64 | 0.28 | <.001 | 19.67 | 0.27 | <.001 |
| Age | 2.49 | 0.04 | <.001 | 3.12 | 0.03 | <.001 |
| Litter size | 0.16 | 0.13 | 0.227 | 0.38 | 0.13 | 0.004 |
| Sex ratio (% male) | 0.54 | 0.15 | <.001 | 0.32 | 0.14 | 0.025 |
| Dom-sired | 0.07 | 0.37 | 0.857 | 1.06 | 0.36 | 0.003 |
| Age x litter size | -0.01 | 0.02 | 0.775 | 0.03 | 0.02 | 0.092 |
| Age x sex ratio | 0.08 | 0.02 | <.001 | 0.04 | 0.02 | 0.027 |
| Age x dom-sired | 0 | 0.05 | 0.989 | 0.18 | 0.04 | <.001 |

Selected outputs from full before-puberty model on round 1 offspring in experiment 2 separated by offspring sex. Paternal factors are shaded in gray and centered in nondominant sired offspring. Age centered at six weeks. Litter size centered at median of seven. Percent of litter that is male is centered at median of 50%. Random effects include animal ID and birthcage. B = coefficient.

### Parental social experience interacts with litter size and sex ratio to regulate body weight

Rodent growth and development is negatively affected by increasing litter size and a male-skewed sex ratio (Drickamer, 1976). We found that pre-puberty growth for daughters and sons of control parents was restricted by 6% and 10% per pup added to a litter, respectively (Fig. 2F; littersize, Table 2). Intriguingly, for competition-sired offspring of all mothers, litter size did not restrict growth (SI tables 2 and 3). Furthermore, dominant-sired daughters and sons of competition mothers actually grow 18% and 15% faster with each additional sibling (Fig. 2F; SI Table 3), suggesting mothers tolerate larger litters if sired by dominant males. In accordance with previous findings on the relationship between sex-ratio and growth rate, control-sired offspring of control mothers grew faster in more male-skewed litters (Table 2). In contrast, dominant-sired offspring of competition mothers grew slower with more males (SI Table 3), while nondominant-sired offspring of competition mothers were unaffected. These results suggest that maternal and paternal effects of social experience can regulate offspring growth by interacting with developmental effects imposed by litter size and sex ratio.

In summary, results from experiment 1 suggest that parental effects of social competition regulate offspring growth throughout development. Notably, paternal social dominance—and to a lesser extent, maternal competition—are drivers of increased growth, and maternal response to nondominant males changes with social experience. Finally, parental effects interact with developmental restrictions like litter size and sex ratio to regulate growth.

## EXPERIMENT 2

### Transitions in paternal social status reversibly affect offspring weight

Results from Experiment 1 suggested that paternal competition promotes offspring growth, and social status modulates this effect. However, although our breeding experiment was designed to statistically uncover parental effects, we were unable to disambiguate direct effects due to dominance status vs. indirect effects of genetic or intrinsic differences between ranks. Indeed, pedigree analysis of experiment 1 males that experienced social competition showed that although dominant and nondominant males were distributed within and between litters, there was a heritable component: maternal birthcage, but not paternal birthcage, was a significant predictor of dominance (binomial logistic regression; paternal birthcage P = 0.649; maternal birthcage P = 0.031; Supplementary Fig. 1B).

To identify plasticity in paternal effects, we designed an experiment to isolate the effects of social dominance transitions while controlling for maternal effects (Fig. 3A, Methods). In the first round, adult males and females (N = 200) on the wild-derived “WLD2” genetic background competed in seminatural enclosures (N = 10) for eight weeks, producing 34 dominant and 44 nondominant males These males were then singly mated in standard cages to a naïve, C57/B6, age-matched female for 10 days, after which the male was removed. These breeding pairs yielded 39 litters and 268 offspring. In the second round, the same males were reintroduced to seminatural enclosures for eight weeks according to their social status: dominant males populated three enclosures and nondominant males populated four enclosures. Dominance was then reassessed, resulting in four paternal conditions: previously dominant males that achieved dominance again (dom->dom, N = 10) or that became nondominant (dom->non, N = 19); and previously nondominant males that remained nondominant (non->non, N = 25) or that gained dominance (non->dom, N = 15). These males were then mated to a new cohort of females, yielding 34 litters and 236 offspring. All available offspring were weighed every week for six weeks (Fig. 3A). We used LMMs and AIC model selection as in experiment 1.

**Figure 3.**
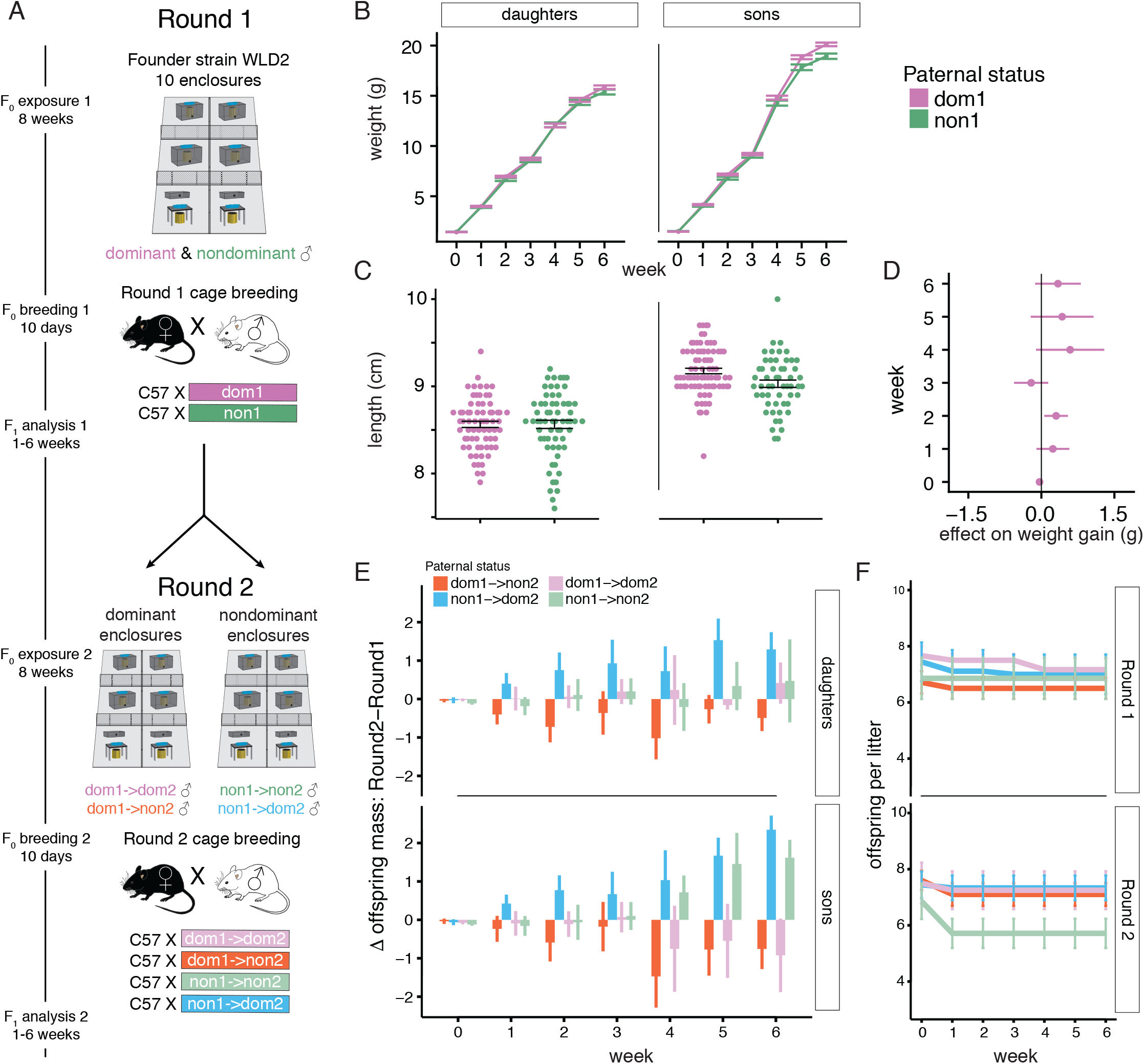
Paternal social status reversibly affects offspring growth. (A) Design of Experiment Two. Round 1: A wild derived strain of mice (WLD2) populated 10 enclosures and male social dominance status was established. After eight weeks males were removed from enclosures and individually assigned to a naive C57B/6 female in a monogamous mating cage (N = 39 litters). Round 2: With naive WLD2 females, dominant males from Round 1 populated three enclosures and nondominant males populated four separate enclosures. After eight weeks all males were categorized as switching or maintaining dominance (i.e., dom1->dom2; dom1->non2; non1->dom2; non1->non2) and assigned to a naïve C57B/6 female in a monogamous mating cage (N = 34 litters). (B-D) Round 1 analysis of sons (N = 134) and daughters (N = 134). (B) Paternal social dominance increases average weight of sons but not daughters during first six weeks of life; mean and SEM. (C) Paternal social dominance increases mean body length in sons but not daughters at six weeks; mean and SEM. (D) Forest plot shows effect of paternal dominance on weight gain in sons while controlling for parental weight and litter size. At each time point symbols and 95% confidence intervals indicate average effect size due to paternal social dominance status relative to nondominant sires (zero line). (E-F) Round 2 analysis of sons (N = 123) and daughters (N = 113). (E) Paternal social dominance reversibly affects offspring weight. Shown is the per-sire mean change (and SEM) in offspring mass from round 1 to round 2 in sons and daughters, error bars that do not cross the zero line represent a significant difference (P < 0.05). Dom1->non2 males produced lighter, whereas non1->dom2 males produced heavier sons and daughters from the first to second round. Non1->non2 males also produced heavier sons. (F) Non->non males show a reduction in litter size during round 2 due to maternal infanticide; shown is mean litter size (and SEM) in round 1 (top) and round 2 (bottom).

After the first round of competition, the full model was the best fit for predicting the weight of sons but not daughters, indicating that paternal status, age, litter size and litter sex ratio collectively regulate the weight of sons (Table 1). Dominant-sired sons grew on average 5% faster than nondominant-sired sons (Table 3), and by six weeks of age were 5% heavier (Fig. 3B, Table 3 Sons) and longer (Fig. 3C; LMM week 6 length, sons: nondominant 9.03 ± 0.31, dominant 9.18 ± 0.28 cm, P < 0.01) than non-dominant sired sons. The effect size of paternal dominance on weekly weight gain was significant at two weeks of age (Fig. 3D). These results confirm that paternal social dominance exerts a growth promoting effect on sons.

We next analyzed how ascending or descending social dominance in the second round of competition affected offspring weight relative to the first round for each father (i.e., intra-individual comparisons). Specifically, we calculated the difference in average offspring weight per litter (for sons and daughters separately) between round two and round one (i.e., R2R1delta, Fig. 3E). Intriguingly, status-descending fathers who went from dominant to nondominant (dom->non) now produced lighter sons (significant at weeks 2, 4, 5 & 6) and lighter daughters (significant at weeks 1, 2, 4 & 6) (Fig. 3E and Supplementary Fig. 2). Status-ascending fathers that went from nondominant to dominant (non->dom) now produced heavier sons and daughters (significant at weeks 1 to 6, Fig. 3E and Supplementary Fig. 2). We then addressed whether the R2R1delta of ascending fathers was different from descending fathers (i.e., inter-individual comparison). Strikingly, the gain in offspring weight for ascending males was significantly greater than the loss in offspring weight for descending males (six week old sons: dom-> non = −0.75 grams; non -> dom = +2.34 grams, P < 0.05; Fig. 3E). There were no differences in between-condition comparisons of males that maintained status across the two rounds (Fig 3E).

Male offspring of doubly-nondominant males exhibited a positive R2R1delta (Fig 3E), an unexpected result because nondominant males typically produce relatively lightweight offspring (Fig. 2 and Fig. 3 B-D). To investigate this further, we evaluated infanticide in round two as a possible mechanism to increase the weight of surviving offspring. Round two cases of infanticide were mostly observed in litters with a nondominant father, and over half (9/17) were in litters with a doubly-nondominant father (Fig. 3F). Accordingly, paternal social status was a significant predictor of the presence/absence of infanticide (binomial logistic regression, effect of paternal status P < 0.001), with doubly nondominant being the most predictive (17.63%). Although offspring from doubly dominant infanticidal litters were heavier than those from non-infanticidal litters at six weeks of age (infanticidal 19.21 ± 0.411, non-infanticidal 18.74 ± 0.839) we lacked statistical power to detect a difference.

When viewed in relation to experiment 1, these results suggest that dominant males produce heavier offspring because of status related cues (not genetic associations); the effect size, timing, and sex-specificity of this paternal effect are modulated by maternal experience. In both experiments, females only had contact with the males for the 10-day breeding opportunity. In experiment 1, the effect of paternal dominance was more pronounced, detected earlier (i.e., week one), and present in both sons and daughters when wild-derived mothers had experienced competition. By contrast, all mothers in experiment 2 were socially naïve C57/B6 females, and here the effect of paternal dominance was detected later (i.e., week two in round one) and in sons but not daughters. Thus, the primary effect of paternal status-dependent regulation of postnatal development of sons can be modulated by mothers to include earlier timepoints and daughters. Doubly nondominant status also affected the likelihood of postnatal maternal infanticide in experiment 2.

### Paternal effects of social status on the liver transcriptome

Results from Experiment 2 confirmed paternal social status induces a nongenetic effect on offspring weight and we sought to identify metabolic pathways and differentially expressed genes (DEG) associated with this effect. Using Bayesian linear regression models, we analyzed pooled RNA from the livers of six-week-old sons from experiment 2 (Methods). For this analysis, offspring were classified by their father’s social status (“D” dominant or “N” nondominant) over the two rounds of competition using the following terminology: DD1, DD2, DN1, DN2, ND1, ND2, NN1, and NN2, where the first letter is round one status, the second is round two status, and the number describes which round the offspring were conceived after. Thus DD2, DN2, ND2, and NN2 describe paternal condition when mating after two rounds of competition. DD1, DS1, SD1, and SS1 describe paternal condition when mating after round one and future round two status.

We first tested the general effect of paternal status (i.e., dominant or nondominant) on liver transcription (Fig. 4A). Several genes were differentially affected by paternal status (SI Table 4.1). In the pooled RNA samples, paternal identity is shared across the two rounds for each condition (e.g., DD1 and DD2 are the same set of fathers, DN1 and DN2 are the same set of fathers, etc.), and we wondered whether paternal identity or social status was a better predictor of offspring transcription. Nonlinear dimensionality reduction using t-SNE (van der Maaten & Hinton, 2008) showed that variation in whole-genome liver transcription is largely explained by paternal status, not paternal identity; sons of nondominant fathers formed one cluster, and sons of dominant fathers formed a separate two clusters (t-SNE1; Fig. 4B). As expected, liver transcription was highly correlated among all offspring, but the sons of doubly-dominant (DD2) and doubly-nondominant (NN2) fathers exhibited the greatest divergence in transcription (Fig. 4C). Gene set enrichment analysis using the Hallmark gene set (Liberzon *et al.*, 2015) showed that DEGs according to paternal status were enriched in pathways relevant to energy metabolism and catabolism, including oxidative phosphorylation, xenobiotic metabolism, and fatty acid metabolism; also detected were changes in mitotic spindle, coagulation, DNA repair, and G2M checkpoint gene sets (FDR adjusted Q < 0.05, Fig 4D).

**Figure 4.**
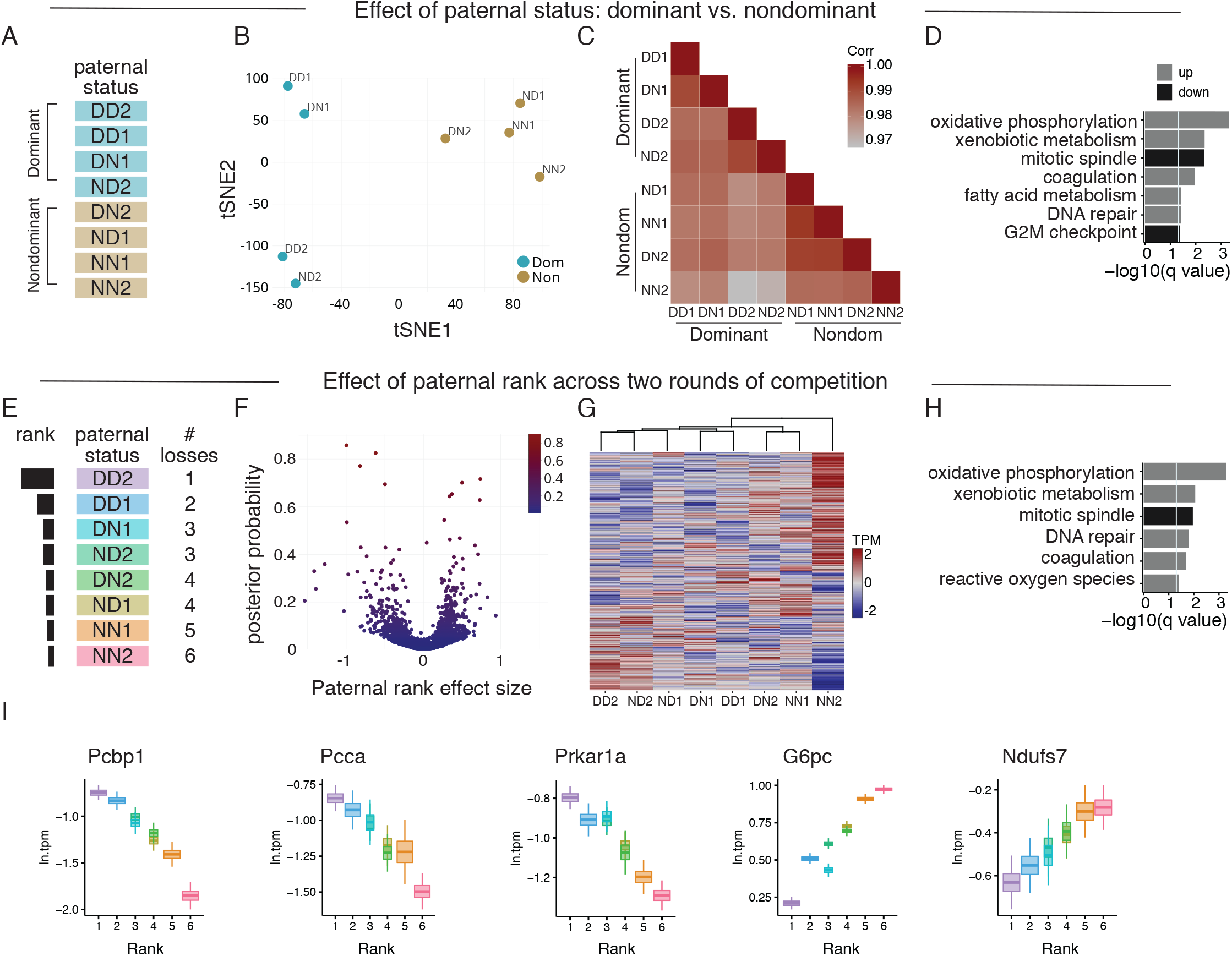
Paternal effects of social status on liver transcription in sons. (A-D) Effect of paternal dominance status on liver transcriptome. Sons (N = 88) were classified according to their father’s dominance status across two rounds of competition and liver RNA was pooled (N = 8 pools; see Methods). (A) Model design; fathers we classified as dominant or nondominant, irrespective of round. (B) tSNE embedding of liver transcription; each point represents sons of the eight paternal conditions and are colored according to dominance status. (C) Correlation matrix of liver transcription according to paternal status. (D) Gene set enrichment analysis (GSEA) of the Hallmark gene sets. Gene sets significantly affected by paternal dominance are shown. The –log(10) FDR-adjusted P value is reported. Vertical line denotes significance threshold –log10(q = 0.05). (E-I) Effect of paternal rank on liver transcriptome across two rounds of competition. (E) Model design; fathers we classified by their hierarchy rank. (F) Volcano plot shows genes according to paternal rank effect size and posterior probability. (G) Hierarchical clustering of liver transcription according to paternal rank. (H) GSEA of Hallmark gene sets affected by paternal rank; values reported as in (D). (I) Candidate genes regulated by paternal rank. Box plots: posterior distribution is the mid line in the box; the top and bottom of the boxes are mean and standard deviation of the posterior distribution; the whiskers are the mean with 3 × standard deviation of the posterior distribution.

We next tested the more specific effect of paternal hierarchy rank on liver transcription (Fig. 4E). To assign hierarchy rank to fathers (and their sons), we devised a metric to score the relative rank for each paternal condition (rank in parentheses): DD2 (1), DD1 (2), DN1 (3), ND2 (3), DN2 (4), ND1 (4), NN1 (5), NN2 (6). Thus, sons with doubly-dominant (DD2) paternity ranked highest, doubly-nondominant (NN2) paternity ranked lowest, and sons with status-switching paternity had an intermediate rank. This analysis yielded several DEGs according to paternal rank (Fig. 4F), several of which were shared with the first analysis (SI Table 4.2). Hierarchical clustering also showed that liver transcription was affected by paternal social rank, forming three clusters (Fig. 4G): cluster-1 was composed of mostly dominant-sired offspring (DD2, ND2, DN1, DD1, but also ND1); cluster-2 was composed of nondominant fathers (DN2 and NN1); cluster-3 was composed of doubly nondominant fathers (NN2) and was more closely associated with cluster-2 than cluster-1. Doubly dominant and doubly nondominant fathers had the most divergent transcriptomes (Fig. 4G). Similarly, t-SNE showed that liver transcription clustered according to paternal rank, not paternal identity (Supplementary Fig. 3A). Gene set enrichment analysis (GSEA) confirmed that several metabolic processes were affected by paternal rank: oxidative phosphorylation, xenobiotic metabolism, and reactive oxygen species; also detected were changes in mitotic spindle, DNA repair, and coagulation (FDR adjusted Q < 0.05, Fig 4H).

Evaluation of the top differentially expressed genes confirmed an association between paternal social rank and metabolic functions (Fig. 4I and Supplementary Fig.3B). Poly(rC) binding protein 1 (*Pcbp1*; posterior probability (PP) = 0.86), is involved in iron metabolism through its delivery of iron to ferritin, an iron storage protein (Nandal *et al.*, 2011), and has been shown to be associated with body weight (Ghanem *et al.*, 2016). Propionyl-Coenzyme A carboxylase alpha polypeptide (*Pcca*; PP = 0.83) is involved in the citric acid cycle and metabolism of branched chain amino acids, odd-numbered chain length fatty acids, and cholesterol; *Pcca* mutations disrupt this process and cause propionic acidemia (Ugarte *et al.*, 1999; Wongkittichote *et al.*, 2017). Two genes involved in glucose metabolism were also identified: *G6pc* and *Prkar1a*. Glucose-6-phosphatase catalytic subunit (*G6pc*, PP = 0.72) plays a primary role in gluconeogenesis by regulating glucose efflux from the cell, and is a causal factor in glycogen storage disease (Mutel *et al.*, 2011). *Prkar1a* (PP = 0.70) is involved with downstream processes of glucagon activated cAMP protein kinase A signaling and is a regulator of the gluconeogenic program (Song *et al.*, 2014). The nuclear-encoded mitochondrial gene NADH:Ubiquinone Oxidoreductase Core Subunit S7 (*Ndufs7*, PP = 0.66) is a subunit of the mitochondrial membrane respiratory chain and is involved with energy metabolism (Mimaki *et al.*, 2012). Additional genes in the top ten were involved in ribosomal processing: *Hnrnpa3* (PP = 0.77); *Nop10*, (PP = 0.70); *Rpl14* (PP = 0.55).

In summary, experiment 2 shows that when controlling for maternal experience, dominant fathers produce heavier sons than nondominant fathers. In addition, status-ascending males produce heavier sons and daughters after gaining dominance, and status-descending males produce lighter sons and daughters after losing dominance. Paternal social rank is also associated with genome-wide liver transcription and metabolic pathways in sons.

## DISCUSSION

Our study on parental effects of social experience on offspring growth has shown that paternal dominance status promotes offspring growth and modulates liver function. In experiment 1, we found that this paternal effect is regulated by maternal experience. Compared to control mothers, competition mothers responded more strongly to the social rank of the father: they produced heavier offspring with dominant males and lighter offspring with nondominant males. In experiment 2 we investigated this paternal effect by using different strains of mice, controlling for maternal experience, and measuring the effect of paternal dominance reversals. Strikingly, ascending males produced heavier sons than before, and descending males produced lighter sons than before. These transitions were also associated with genome-wide hepatic transcription in sons.

The reversibility of parental effects is an important, but understudied feature of adaptation. Parental effects can persist after the removal of environmental exposures for several generations, but in a fluctuating environment it is crucial they happen on a timescale commensurate with the breeding cycle—if not, they could be maladaptive (Gabriel *et al.*, 2005). Social dominance hierarchies take days to weeks to be established (Nelson *et al.*, 2013b; Nelson *et al.*, 2015), and we find that signals about paternal rank can be transmitted to offspring on this timescale. Previously, we found that dominant males have lower locomotor economy than nondominant males, suggesting a tradeoff between fighting ability and locomotion (Morris *et al.*, 2017). Thus, promoting fast-growing and powerful sons could be adaptive under conditions of elevated male-male competition (e.g., in a dominant territory). Conversely, promoting slow growing sons with higher locomotor economy could be adaptive under conditions of predation, resource limitation and dispersal (e.g., in areas inhabited by nondominants).

Stress and nutritional manipulations have shown that parental environment can regulate offspring metabolic phenotype (Rando, 2012; Rando & Simmons, 2015). Our finding that paternal dominance status regulates offspring liver transcription provides context for how these metabolic processes might occur in natural situations. Parental reprogramming of mitochondrial function appears to be a key transgenerational mediator of metabolic phenotype (Rando & Simmons, 2015). Accordingly, we identified two nuclear-mitochondrial genes that were affected by paternal rank. *Pcca* plays a role in the citric acid cycle through synthesis of succinyl-CoA (Wongkittichote *et al.*, 2017), and *Ndfus7* is involved with energy metabolism (Mimaki *et al.*, 2012). Glucose homeostasis is also subject to considerable regulation by maternal and paternal environment (Rando & Simmons, 2015). We identified two genes involved in glucose metabolism: *G6pc* and *Prkar1a*, both of which are involved in gluconeogenesis (Mutel *et al.*, 2011; Song *et al.*, 2014). Our GSEA findings are consistent with several other studies that found paternal effects on sugar, lipid, and xenobiotic metabolism, and emphasize the possibility that rodent paternal effects feed into a few pleiotropic offspring response programs that mediate tradeoffs between stress and growth (Rando & Simmons, 2015; Vallaster *et al.*, 2017).

This study provides of point of entry for dissecting the mechanistic basis of paternal effects of social status. At the organismal level, there are two non-exclusive hypotheses to explain our results: (1) maternal interpretation of paternal quality or (2) a direct paternal effect delivered through sperm. In the first scenario, maternal interpretation of paternal quality results in differential allocation of investment to developing offspring either prenatally or postnatally (Burley, 1988; Cunningham & Russell, 2000; Mashoodh *et al.*, 2012). In the second scenario, information about paternal condition is transferred from the father to the mother via factors in the seminal fluid or epigenetic modifications of sperm (Watkins *et al.*, 2018). These two hypotheses could be addressed with cross-fostering, in vitro fertilization, artificial insemination, and embryo transfer experiments. At the molecular level, determining the causal relationships between the metabolic gene expression and body mass will be an essential step forward.

## Acknowledgements

We thank Brad Cairns for stimulating discussions, Linda Morrison for colony management and Yajaira Peralta and Derek Stark for assisting with the management of semi-natural enclosures and data collection. This work was supported while W.K.P was supported by NIH grant R01-GM109500, in conjunction with the NSF-managed Ecology and Evolution of Infectious Disease (EEID) program. L.N.S and J.A.B. were supported by the Undergraduate Research Opportunities Program at the University of Utah. Yajaira Peralta was supported by the NSF funded Western Alliance to Expand Student Opportunities (WAESO).

## MAJOR METHODS

### ANIMALS

#### Experiment 1 animals

Mice used in experiment 1 were derived from a cross between wild-caught mice and MHC-congenic mice carrying five known haplotypes (C57BL/10SnJ-H2b, B10.D2-H2d, B10.M-H2f, B10.BR-H2k, and B10.Q-H2q) obtained from The Jackson Laboratory, where wild-derived MHC haplotypes were eliminated by selective breeding (Ilmonen *et al.*, 2007). This “congenic/wild” or “CNGWLD” strain has been bred in the Department of Biology at University of Utah for over 10 generations, and has been used to study naturalistic social behavior (Nelson *et al.*, 2013a; Nelson *et al.*, 2013b). To determine parental effects of social experience on offspring weight, 168 adult mice (12 to 20 weeks of age) were randomly assigned to either an eight-week competition experience in semi-natural enclosures or an eight-week monogamous breeding cage experience. In most cases, siblings were split between the two conditions. In competition groups, males were assessed for territoriality and classified as either dominant or nondominant and females were classified as competition. In monogamous pairs, males and females were classified as control. After the eight-week exposure, half of the competition females were removed from the enclosures and all monogamous breeding cages were separated for three weeks (this step insured that all females were not pregnant when entering the reciprocal breeding experiment). During this three-week interval, the other half of the competition females were left in the enclosures to help continue normal male territoriality. Following this three-week interval, a reciprocal breeding design was employed to systematically assess parent of origin effects of social experience and status on offspring body mass. Specifically, all combinations of males (dominant, nondominant, control) and females (competition, control) were bred as monogamous pairs in standard cages for 10 days, after which the male was removed. These pairs yielded 56 litters total (420 offspring; 198 daughters and 222 sons); 7 Control mother x Dominant father litters (54 offspring; 25 daughters and 29 sons), 7 Control x Nondominant (62 offs; 27 daughters and 35 sons), 10 Control x Control (71 offs; 31 daughters and 40 sons), 6 Competition x Dominant (42 offs; 17 daughters and 25 sons), 4 Competition x Nondominant (32 offs; 17 daughters and 15 sons), and 22 Competition x Control (159 offs; 81 daughters, 78 sons). The resulting offspring were then assessed for growth rate.

#### Experiment 2 animals

Mice used in experiment 2 were from an outbred, wild-derived “WLD2” strain bred for 18 generations at the University of Utah and previously described (Meagher *et al.*, 2000). Genetic diversity was assessed during the 11^th^ generation and found to be comparable to wild populations (Cunningham *et al.*, 2013). Experiment 2 resembles experiment 1 in several ways, but differs in that all offspring had C57/B6 maternal chromosomes, and wild-derived paternal chromosomes, as opposed to an average half-and-half background. To target paternal effects of maintaining or losing social status, experiment 2 proceeded across two rounds of competition. In the first round, 100 male and 100 female founders competed in 10 semi-natural enclosures for eight weeks. These enclosures produced 34 dominant males and 44 nondominant males (22 died or were euthanized due to injury, ~10-20% mortality is expected). The 78 males were then singly mated in standard cages to a naïve, C57/B6, age-matched (8 weeks old) female for 10 days, after which the male was removed. These breeding pairs yielded 39 litters total (268 offspring; 134 daughters and 134 sons): 22 from dominant fathers (150 offs; 70 daughters and 80 sons) and 17 from non-dominant fathers (118 offs; 64 daughters and 54 sons). In the second round, the males were reintroduced to semi-natural enclosures for eight weeks according to their social status; dominant males populated three enclosures and nondominant males populated four enclosures. Each enclosure contained 11 males and 11 females. Dominance was reassessed for the 69 males that completed the second round of competition (8 males died during competition); 10 previously dominant males achieved dominance again (DD), while 19 failed to achieve dominance a second time (DN); 15 previously non-dominant males newly achieved dominance (ND), while 25 failed a second time (NN). These males were then singly mated to a new cohort of age-matched C57/B6 females for 10 days, after which the males were removed and sacrificed for tissue collection. These breeding pairs yielded 34 litters total (236 offspring; 113 daughters and 123 sons): 8 from Dom->Dom (58 offs; 29 daughters and 29 sons), 10 Dom->Non (71 offs; 29 daughters and 42 sons), 9 Non->Dom (67 offs; 36 daughters and 31 sons), and 7 Non->Non (40 offs; 19 daughters and 21 sons). During both rounds of breeding, females remained individually housed after the removal of the male and monitored for significant weight gain at day 19 to predict parturition. Pup checks were performed every 12 hours, allowing all pups to have their birth mass and birthday accurately recorded. To make repeated measures pups were uniquely marked by toe-clipping (elaborated in “toe-clipping”). Pups were weighed every seven days. At exactly six weeks of age, pups were dissected to collect whole livers. All animal methods and practices are approval by the IACUC at University of Utah these experiments.

#### Experiment 2 social status nomenclature

Offspring conceived in experiment 2 were classified by their fathers’ social status (“D” dominant or “N” nondominant) over the two rounds, producing the following nomenclature: DD1, DD2, DN1, DN2, ND1, ND2, NN1, and NN2. The first letter is round one status, the second is round two status, and tailing number describes which round the offspring were conceived after. So DD2, DN2, ND2, and NN2 describe the total social history for each father up to the point of conceiving those litters, while information about the father’s future is included for DD1, DS1, SD1, and SS1, because the second rounds had not yet occurred. To assign hierarchy rank to the eight paternal categories, we scored the number of competitive losses experienced by each group (losses in parentheses): DD2 (1), DD1 (2), DN1 (3), ND2 (3), DN2 (4), ND1 (4), NN1 (5), NN2 (6). In this scheme, DD1 had the fewest losses and highest rank, NN2 had the most losses and lowest rank, and males that switched status had an intermediate number of losses.

### SOCIAL DOMINANCE AND BODY MASS

#### Semi-natural enclosures

Enclosures were ~30 m^2^ in area and subdivided into six sub-sections with food and water sources provided *ad libitum*. “Optimal” sub-sections contained an enclosed nest box containing food (**Fig. 1**: territories 2-5) and “suboptimal” sub-sections contained an exposed nest box that was separate from food (**Fig. 1:** territories 1 and 6). This layout has been developed to trigger competition over breeding sites and typically produces clear dominant males and non-dominant males. On average, dominant mice occupy the four optimal territories whereas the two suboptimal territories contain nondominant individuals who would likely disperse in a natural setting. Enclosures were founded by 10 males and 10 females unless otherwise noted.

#### Social dominance evaluation

All animals, including those designated for breeding cage experience, were implanted with passive integrated transponder (PIT) tags. Tags in competition animals were used to aid in identification and quantification of social dominance and survival. A male is determined dominant if he possesses more than 80% of total male PIT tag reads in a given territory (Nelson *et al.*, 2013a; Nelson *et al.*, 2013b). Female territoriality is also present in these enclosures but is not as obvious to an observer or as quantifiable with pit-tag data.

#### Toe-clipping

Offspring in experiment 1 and 2 were uniquely marked using a system based on NIH toe-clipping guidelines. Briefly, a sharp sterile pair of scissors were used to, at most, amputate the third segment of one non-dew claw or index toe on one paw (three toes on each paw possible), prioritizing hind paws. This allowed a numbering system to be designed that counted up to 13 (3 toes x 4 paws + no clip). To achieve a unique mark, the minimal amount needed to be removed from a toe was the nail bed, as identifying which toe had been modified was consistent and complete by observing the absence of a toenail.

#### Body mass measurements

All body mass data was collected by briefly placing the animal into a clear plastic zip-lock bag, then on to a scale accurate to the hundredth of a gram.

#### Experiment 1

F0 (founders) males and females were weighed at multiple time-points: one day prior to social experience during handling for pit-tagging (10-12 weeks of age), body-mass at treatment was measure during the 8 week pup sweep (18-20 weeks old), and body mass at sacrifice (19-21 weeks old). No female founders were sacrificed following treatment in order to mate, gestate, and give birth to F1s. Experiment 1 offspring were weighed and toe-clipped at one week old, then repeatedly weighed at 3 weeks (weaning), 5, 6, 8, 10, 12, 14, 16, 18, 20, 22, 24, 26, 28, and 30 weeks of age, and females were additionally measured at 32 weeks. All age points are the body mass of the animals at precisely “x” weeks from birth, accurate up to 12 hours after birth. If the precise day was missed, those data were not recorded. Male offspring were individually housed at 22 weeks old to standardize social environment and sacrificed at 30 weeks; therefore, no male offspring have a recorded 32-week body-mass.

#### Experiment 2

Male founders were weighed before and after each round of competition, and at sacrifice. Breeder females were weighed before their breeding opportunity and during gestation. All experiment 2 offspring were weighed within 12 hours of birth and every 7 days until six week of age. All animals were sacrificed and dissected within 12 hours of being six weeks old.

### RNA-SEQ

#### Founder and offspring liver dissection

All animals were euthanized with CO_2_ and secondarily euthanized by cervical dislocation. Same sex siblings were sacrificed together and dissected quickly in order of toe-clips. Each dissection took about 5-7 minutes and was performed by two technicians. Dissections began by pinning and opening the animal ventrally from genitals to diaphragm. Next, the entire liver is removed, positioned with median lobe external, wrapped in labeled tinfoil, and flash frozen in liquid nitrogen for storage until RNA isolation.

#### RNA isolation

liver RNA was extracted from offspring after both rounds of experiment 2. Two sons from each round for each father (four sons total) were selected favoring the earliest dissections for each litter (lowest toe-clip). 88 sons, ^~^11 from each type, were extracted using a standard TRIzol protocol (Gapp *et al.*, 2014; Sharma *et al.*, 2016). Briefly, a ^~^20-50 mg piece of liver was removed from the median lobe of a still frozen whole liver (using dry ice and a cold room), and placed in an Eppendorf tube with 1 mL of TRIzol. The liver chuck was then completely pulverized using a tight-fit plastic homogenizer. Once homogenous, 200 uL of chloroform was added and incubated at room temperature for 3 minutes. Samples were then spun cold (4°C) at 12000 G for 15 minutes. The aqueous phase (^~^500 uL) was then transferred to a new tube and the RNA was precipitated out by adding 500 uL isopropanol and similarly spinning cold at 12000 G for 15 minutes. RNA pellets were then washed in 1 mL of 75% ethanol, and re-suspended in 50 ul of RNase-free water.

#### RNA-Seq, mapping, and expression estimation

Eight pools of son RNA, all from fathers with offspring in both rounds, were constructed for sequencing. Each son contributed approximately equal (20 ng total RNA) amounts of RNA to each pool, estimated using Nano Drop (Nano Drop Technologies, San Diego, CA) report concentrations. The eight pools were classified according to experiment 2 nomenclature described above (DD1, DN1, ND1, NN1, DD2, DN2, ND2, NN2). Strand-specific libraries were prepared as previously described (Zhang *et al.*, 2012). After first-and second-strand synthesis, adapters were ligated to fragments and amplified using multiplexed PCR primers. Libraries were sequenced on a NextSeq 500 (Illumina Inc., San Diego, CA, USA), generating a range of 26,574,081-46,832,331 79 bp paired-end reads.

RNA-seq library mapping and estimation of expression levels were computed as follows. Reads were mapped with STAR aligner version 2.5.3a (Dobin *et al.*, 2013), using the two-round mapping approach, to the mm10 reference mouse genome and the Gencode vM12 primary assembly annotation (Mudge & Harrow, 2015), to which non-redundant UCSC transcripts were added. This means that following a first mapping round of each library, the splice-junction coordinates reported by STAR, across all libraries, were fed as input to the second round of mapping. The parameters used in both mapping rounds were: outSAMprimaryFlag = “AllBestScore”, outFilterMultimapNmax = “10”, outFilterMismatchNoverLmax = “0.05”, outFilterIntronMotifs = “RemoveNoncanonical”. Following read mapping, transcript and gene expression levels were estimated using MMSEQ (Turro *et al.*, 2011). MMSEQ employs a Bayesian model for estimating transcript-level expression and hence reports the posterior distribution of these expression estimates. Next, transcripts and genes which could not be distinguished according to read data were collapsed using the mmcollapse utility of MMDIFF (Turro *et al.*, 2014). Fragments Per Killobase Million (FPKM) expression units were converted to Transcript Per Million (TPM) units since the latter were shown to be less biased and more interpretable (Wagner *et al.*, 2012).

### STATISTICAL ANALYSES

#### Body mass and growth rate analysis

Growth rates of the offspring were compared using linear mixed models (LMMs). Previously reported predictors of body mass were included: time (age), sex, litter size, and sex ratio. Paternal dominance and maternal treatment (experiment 1) were included as our tested factors. These factors and all possible interactions were modeled as fixed effects while parents’ populations, parent litter IDs, offspring IDs, and offspring litter IDs are modeled as random effects. The random effects of parents’ population, and parents’ litter, which accounted for variation between enclosures and cousin level relatedness of parents, did not contribute meaningfully to the model The random effects that remained controlled for pseudo-replication of siblings within each litter (litter IDs), and repeated measures across time (offspring ID). Because offspring grow at different, but near linear, rates before and after puberty (i.e., six weeks of age), and because parental effects were different before and after puberty, we analyzed these two developmental stages using separate LMMs. The best fit model was selected from multiple rounds of systematic fixed effect, interaction, and random effect elimination and re-introduction. The lowest AICc, by greater than 2, was favored as best fit. LMMs were performed in R using the lme4 library (R-Development-Core-Team, 2011; Bates *et al.*, 2015). To test hypotheses about categorical dependent variables (i.e., heritability of dominance in experiment 1, and presence/absence of infanticide in experiment 2), we used binomial logistic regression in R.

#### RNA-seq analysis

To estimate the effect of birth status on ln(TPM) units of gene expression (i.e., differential expression between dominant and subordinate) we used MMDIFF (Turro *et al.*, 2014), with the following model:

M: 0 0 0 0 0 0 0 0. C: 0 0, 0 0, 0 0, 0 0, 0 1, 0 1, 0 1, 0 1. Where the samples are in order: DD2, DD1, DS1, SD2, DS2, SD1, SS1, and SS2, corresponding to MMDIFF M and C matrices. To estimate the effect of hierarchy rank (i.e. number of losses) on ln(TPM), we used also used MMDIFF, with the following model: M: 0 0 0 0 0 0 0 0. C: 0 0 0, 1 0, 2 0, 2 0, 3 0, 3 0, 4 0, 5 0. P0: 1. P1: 0 0.2 0.4 0.6 0.8 1. Where the samples are in the same order as for the birth status model: DD2, DD1, DS1, SD2, DS2, SD1, SS1, and SS2, corresponding to MMDIFF M and C matrices.

In both analyses we empirically determined the limit of detection to be at the main mode of the distribution of gene mean ln(TPM). For analysis of birth status the mean and limit of detection were computed separately for each of the two birth status categories and found to be −1.9 ln(TPM) for the subordinates and −2.1 for the dominants. For analysis of hierarchy rank, the mean ln(TPM) was taken across all samples and hence the limit of detection was found to be −2.2 ln(TPM). Thus, in each analysis, any gene with an ln(TPM) below its corresponding limit of detection, in any given sample, was set to limit of detection (to avoid taking logs of zero). Finally, in the analysis of hierarchy rank we required a gene to have ln(TPM) values below the limit of detection in at least two samples. Moreover, in order to avoid including genes which expression levels are mainly based on ambiguously mapped reads and may thus be unreliable, we also required a gene to have at least four uniquely mapped reads in at least two samples. Genes which did not meet these requirements were filtered from the analysis. For the analysis of birth status, we took the same filtering approach, but this time required that the conditions above be met for at least two samples among the four samples of each of the birth status categories.

In both analyses we ranked the genes according to the Bayes factor obtained by contrasting a model which assumes that ln(TPM) is explained by the factor of interest (i.e., birth status or social rank) with a null model in which the factor of interest is dropped. In addition, we applied a secondary ranking according to the posterior probability of the effect being different from zero. For the analysis of rank, a significance threshold posterior probability of 0.65 was choses empirically, as it appeared to be the point beyond which the likelihood of an effect different from zero dominated the low prior (0.1) (Supplementary Fig. 3B).

To test for paternal effects of hierarchy status on transcriptional pathways in offspring, we used Gene Set Enrichment Analysis (GSEA, (Sergushichev, 2016)) of the Hallmark gene set obtained from mSigDB collections (Liberzon *et al.*, 2015). The Hallmark sets are composed of genes that display overlapping and coordinated expression in other mSigDB collections, and summarize well-defined biological states or processes. Additional downstream analyses included generating a sample ln(TPM) pairwise Pearson correlation coefficient heatmap, and a genes-by-samples centered and scaled ln(TPM) hierarchically clustered heatmap (using the fastcluster R package (Müllner, 2013)).

---

**Supplementary Figure 1.**
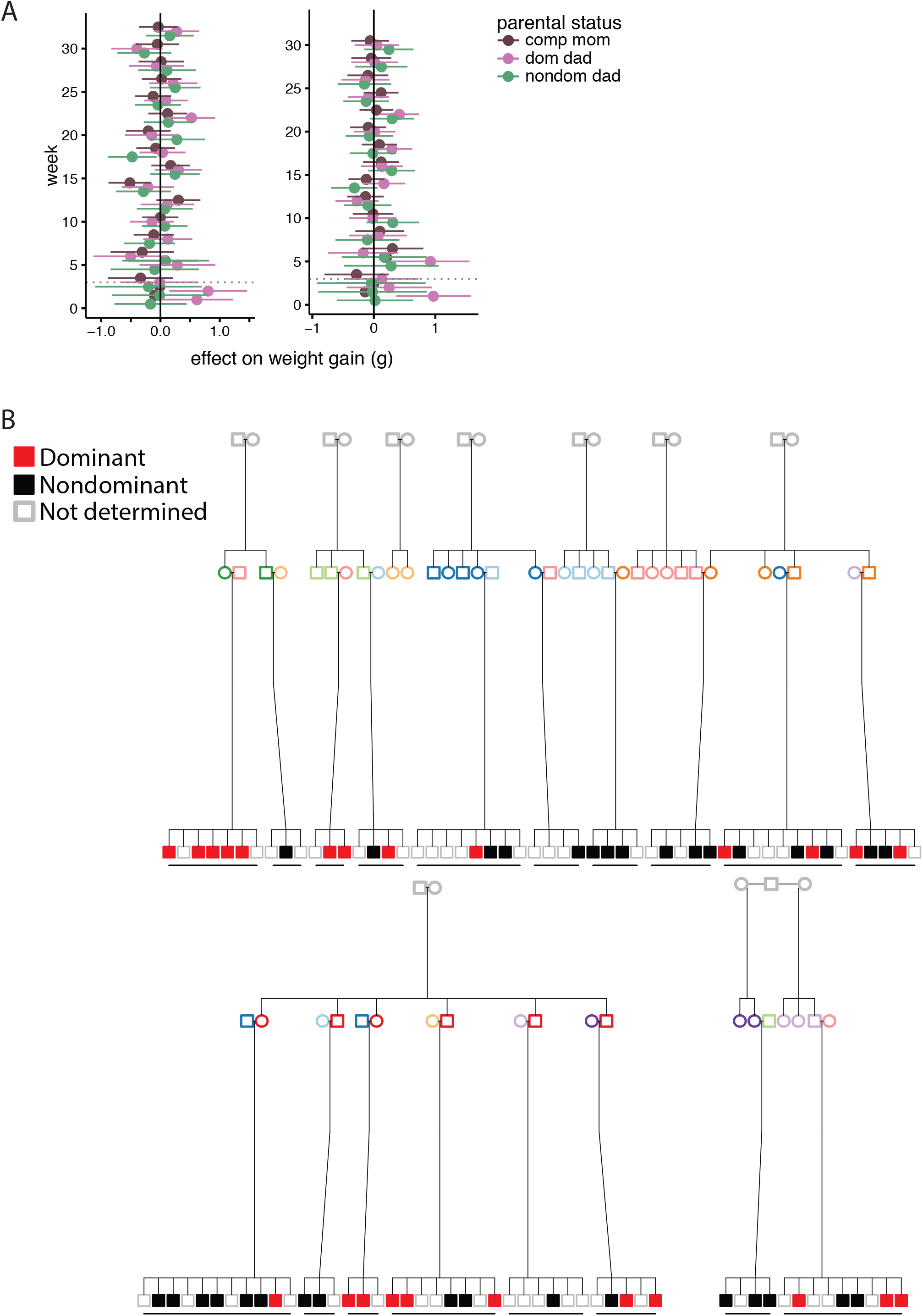
Experiment 1. (A) Forest plots show the association between parental social status and weekly weight gain of daughters (left, N = 113) and sons (right, N = 123) while controlling for other factors; data are coefficients from LMM at each time point separately. Means and 95% confidence intervals indicate the effect size on body weight due to maternal social competition and paternal social dominance status relative to monogamous control parents (zero line). Dotted lines at week 3 indicate time of weaning. (B) Pedigree of males (i.e., fathers) from experiment 1 (N = 18 litters and 101 males). Dominant (red), nondominant (black) and non-determined (grey) males are categorized according to their litter. Mothers (colored circles), fathers (colored squares) and grandparents (gray circles and squares) of these males are also shown. Maternal birthcage, but not paternal birthcage, was a significant predictor of social dominance (see text).

**Supplementary Figure 2.**
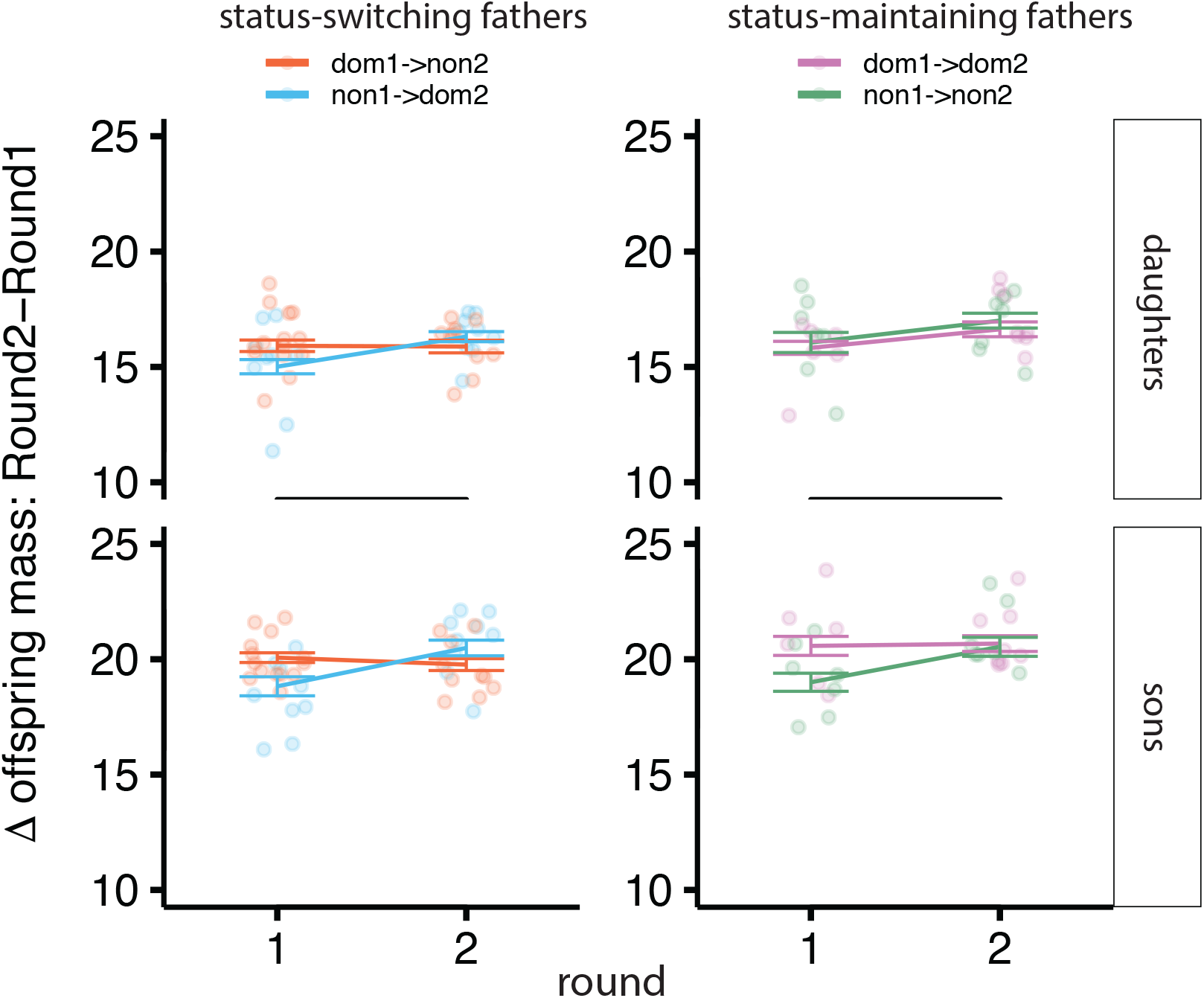
Experiment 2. Change in mean weight of six-week old offspring across two rounds of competition. Left: offspring of status-switching males (N = 19 litters; 10 Dom->Non, 9 Non->Dom). Right: offspring of status-maintaining males (N = 15 litters; 8 Dom->Dom, 7 Non->Non). Each data point is the mean from one litter, lines show means and SEM for each condition.

**Supplementary Figure 3.**
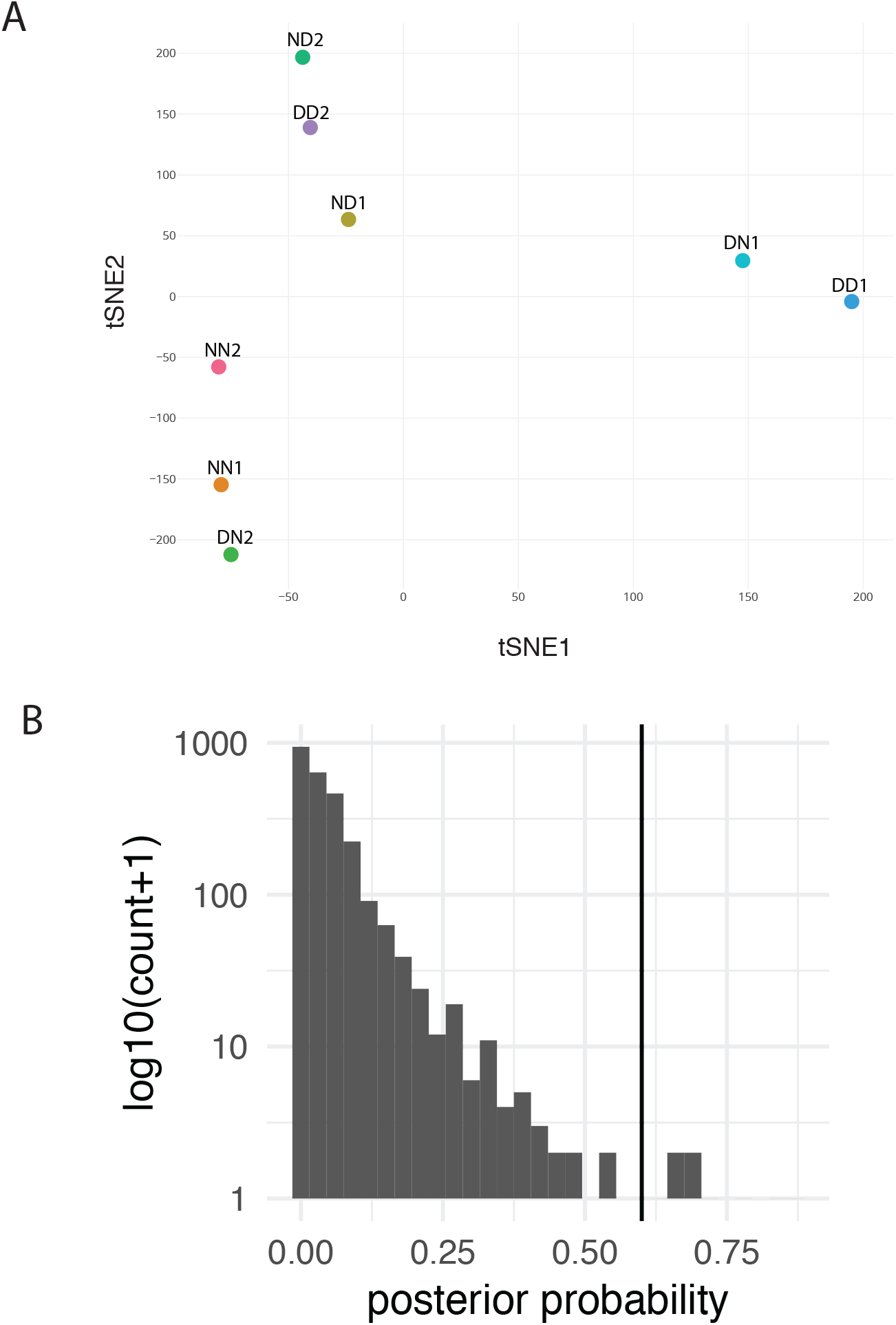
Experiment 2 RNA-seq. (A) Nonlinear dimensionality reduction using t-distributed Stochastic Neighbor Embedding (t-SNE) of liver transcription and colored according to paternal rank (B) Effects significance histogram shows the log distribution of posterior probability values.

**SI Table 1.** Post-puberty predictors of offspring weight in Experiment 1. Refers to Figure 2.

|  | Control mother |  |  | Competition mother |  |  |
| --- | --- | --- | --- | --- | --- | --- |
|  | *B* | *Std. error* | *P* | *B* | *Std. error* | *P* |
| **Daughters** |  |  |  |  |  |  |
| Intercept | 19.28 | 0.73 | <.001 | 20.97 | 0.33 | <.001 |
| Litter size | -0.96 | 0.39 | 0.014 | 0.07 | 0.23 | 0.762 |
| Sex ratio (% male) | 1.06 | 0.53 | 0.045 | -0.05 | 0.19 | 0.775 |
| Dom-sired | 3.08 | 0.91 | <.001 | 3.56 | 1.27 | 0.005 |
| Nondom-sired | 2.01 | 0.99 | 0.043 | -1.81 | 1.27 | 0.155 |
| Maternal condition | -1.68 | 0.80 | 0.35 | 1.68 | 0.80 | 0.35 |
| Age (g/week) | 0.19 | 0.02 | <.001 | 0.21 | 0.01 | <.001 |
| Age x litter size | 0.00 | 0.01 | 0.91 | 0.00 | 0.01 | 0.363 |
| Age x sex ratio | 0.00 | 0.01 | 0.702 | 0.00 | 0.00 | 0.722 |
| Age x dom-sired | 0.05 | 0.02 | 0.021 | 0.06 | 0.03 | 0.022 |
| Age x nondom-sired | 0.03 | 0.02 | 0.242 | -0.05 | 0.03 | 0.063 |
| Age x maternal cond. | -0.01 | 0.02 | 0.492 | 0.01 | 0.02 | 0.492 |
| **Sons** |  |  |  |  |  |  |
| Intercept | 24.08 | 0.72 | <.001 | 27.3 | 0.33 | <.001 |
| Litter size | -0.84 | 0.38 | 0.029 | -0.7 | 0.22 | 0.001 |
| Sex ratio (% male) | 2.05 | 0.44 | <.001 | -0.31 | 0.21 | 0.144 |
| Dom-sired | 6.36 | 0.9 | <.001 | 4.37 | 1.29 | <.001 |
| Nondom-sired | 4.18 | 0.99 | <.001 | -1.26 | 1.29 | 0.331 |
| Maternal condition | 3.22 | 0.79 | <.001 | -3.22 | 0.79 | <.001 |
| Age (g/week) | 0.3 | 0.02 | <.001 | 0.31 | 0.01 | <.001 |
| Age x litter size | 0.01 | 0.01 | 0.209 | -0.01 | 0.01 | 0.107 |
| Age x sex ratio | 0.00 | 0.01 | 0.838 | -0.01 | 0.01 | 0.294 |
| Age x dom-sired | 0.08 | 0.02 | <.001 | 0.03 | 0.03 | 0.393 |
| Age x nondom-sired | 0.02 | 0.02 | 0.43 | -0.1 | 0.03 | 0.002 |
| Age x maternal cond. | -0.01 | 0.02 | .609 | 0.01 | 0.02 | .609 |

Selected outputs from full after-puberty model separated by offspring sex and maternal condition. Left columns are leveled on control mothers and right columns are leveled on competition mothers. Paternal factors are shaded in gray. Age centered at thirty weeks. Litter size centered at median of eight pups. Percent of litter that is male is centered at median of 50%. Parental effects of social experience are in comparison to control-sired offspring. Random effects include animal ID and birthcage. B = coefficient.

**SI Table 2.**
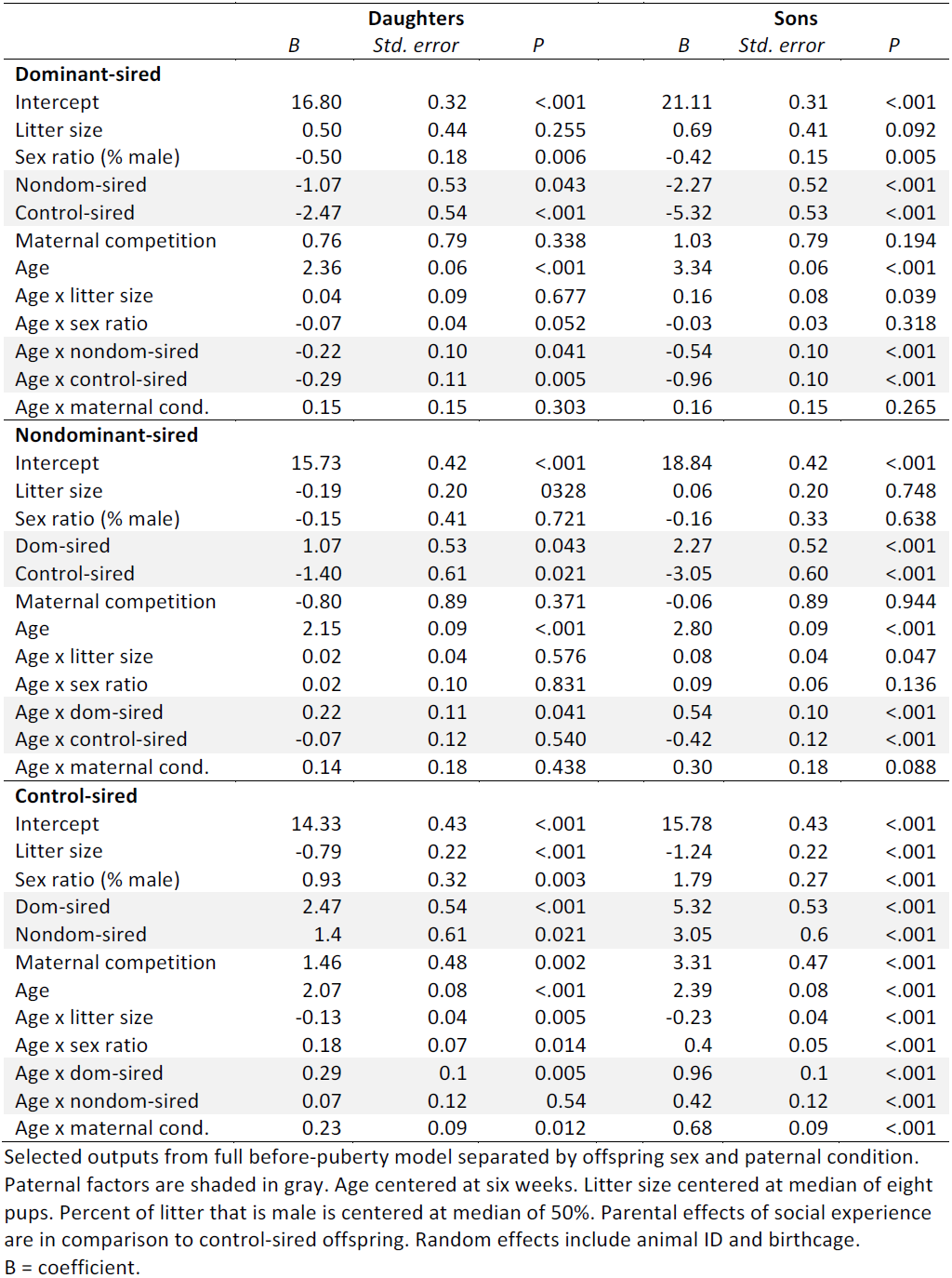
Before puberty predictors of offspring weight of control mothers in Exp. 1 separated by sires’ condition.

|  | **Daughters** |  |  | **Sons** |  |  |
| --- | --- | --- | --- | --- | --- | --- |
|  | *B* | *Std. error* | *P* | *B* | *Std. error* | *P* |
| **Dominant-sired** |  |  |  |  |  |  |
| Intercept | 16.80 | 0.32 | <.001 | 21.11 | 0.31 | <.001 |
| Litter size | 0.50 | 0.44 | 0.255 | 0.69 | 0.41 | 0.092 |
| Sex ratio (% male) | -0.50 | 0.18 | 0.006 | -0.42 | 0.15 | 0.005 |
| Nondom-sired | -1.07 | 0.53 | 0.043 | -2.27 | 0.52 | <.001 |
| Control-sired | -2.47 | 0.54 | <.001 | -5.32 | 0.53 | <.001 |
| Maternal competition | 0.76 | 0.79 | 0.338 | 1.03 | 0.79 | 0.194 |
| Age | 2.36 | 0.06 | <.001 | 3.34 | 0.06 | <.001 |
| Age x litter size | 0.04 | 0.09 | 0.677 | 0.16 | 0.08 | 0.039 |
| Age x sex ratio | -0.07 | 0.04 | 0.052 | -0.03 | 0.03 | 0.318 |
| Age x nondom-sired | -0.22 | 0.10 | 0.041 | -0.54 | 0.10 | <.001 |
| Age x control-sired | -0.29 | 0.11 | 0.005 | -0.96 | 0.10 | <.001 |
| Age x maternal cond. | 0.15 | 0.15 | 0.303 | 0.16 | 0.15 | 0.265 |
| **Nondominant-sired** |  |  |  |  |  |  |
| Intercept | 15.73 | 0.42 | <.001 | 18.84 | 0.42 | <.001 |
| Litter size | -0.19 | 0.20 | 0.328 | 0.06 | 0.20 | 0.748 |
| Sex ratio (% male) | -0.15 | 0.41 | 0.721 | -0.16 | 0.33 | 0.638 |
| Dom-sired | 1.07 | 0.53 | 0.043 | 2.27 | 0.52 | <.001 |
| Control-sired | -1.40 | 0.61 | 0.021 | -3.05 | 0.60 | <.001 |
| Maternal competition | -0.80 | 0.89 | 0.371 | -0.06 | 0.89 | 0.944 |
| Age | 2.15 | 0.09 | <.001 | 2.80 | 0.09 | <.001 |
| Age x litter size | 0.02 | 0.04 | 0.576 | 0.08 | 0.04 | 0.047 |
| Age x sex ratio | 0.02 | 0.10 | 0.831 | 0.09 | 0.06 | 0.136 |
| Age x dom-sired | 0.22 | 0.11 | 0.041 | 0.54 | 0.10 | <.001 |
| Age x control-sired | -0.07 | 0.12 | 0.540 | -0.42 | 0.12 | <.001 |
| Age x maternal cond. | 0.14 | 0.18 | 0.438 | 0.30 | 0.18 | 0.088 |
| **Control-sired** |  |  |  |  |  |  |
| Intercept | 14.33 | 0.43 | <.001 | 15.78 | 0.43 | <.001 |
| Litter size | -0.79 | 0.22 | <.001 | -1.24 | 0.22 | <.001 |
| Sex ratio (% male) | 0.93 | 0.32 | 0.003 | 1.79 | 0.27 | <.001 |
| Dom-sired | 2.47 | 0.54 | <.001 | 5.32 | 0.53 | <.001 |
| Nondom-sired | 1.4 | 0.61 | 0.021 | 3.05 | 0.6 | <.001 |
| Maternal competition | 1.46 | 0.48 | 0.002 | 3.31 | 0.47 | <.001 |
| Age | 2.07 | 0.08 | <.001 | 2.39 | 0.08 | <.001 |
| Age x litter size | -0.13 | 0.04 | 0.005 | -0.23 | 0.04 | <.001 |
| Age x sex ratio | 0.18 | 0.07 | 0.014 | 0.4 | 0.05 | <.001 |
| Age x dom-sired | 0.29 | 0.1 | 0.005 | 0.96 | 0.1 | <.001 |
| Age x nondom-sired | 0.07 | 0.12 | 0.54 | 0.42 | 0.12 | <.001 |
| Age x maternal cond. | 0.23 | 0.09 | 0.012 | 0.68 | 0.09 | <.001 |

Selected outputs from full before-puberty model separated by offspring sex and paternal condition. Paternal factors are shaded in gray. Age centered at six weeks. Litter size centered at median of eight pups. Percent of litter that is male is centered at median of 50%. Parental effects of social experience are in comparison to control-sired offspring. Random effects include animal ID and birthcage. B = coefficient.

**SI Table 3.** Before puberty predictors of offspring weight of competition mothers in Exp. 1 separated by sires’ condition.

|  | Daughters |  |  | Sons |  |  |
| --- | --- | --- | --- | --- | --- | --- |
|  | *B* | *Std. error* | *P* | *B* | *Std. error* | *P* |
| **Dominant-sired** |  |  |  |  |  |  |
| Intercept | 17.56 | 0.73 | <.001 | 22.13 | 0.73 | <.001 |
| Litter size | 3.11 | 0.90 | <.001 | 4.03 | 0.89 | <.001 |
| Sex ratio (% male) | -2.95 | 0.75 | <.001 | -3.14 | 0.49 | <.001 |
| Nondom-sired control | -2.63 | 1.07 | 0.014 | -3.36 | 1.07 | 0.002 |
| Control-sired | -1.77 | 0.75 | 0.019 | -3.04 | 0.75 | <.001 |
| Maternal control | -0.76 | 0.79 | 0.338 | -1.03 | 0.79 | 0.194 |
| Age | 2.51 | 0.13 | <.001 | 3.51 | 0.13 | <.001 |
| Age x litter size | 0.45 | 0.18 | 0.014 | 0.54 | 0.18 | 0.002 |
| Age x sex ratio | -0.37 | 0.20 | 0.061 | -0.31 | 0.10 | 0.002 |
| Age x nondom-sired | -0.23 | 0.20 | 0.260 | -0.40 | 0.20 | 0.047 |
| Age x control-sired | -0.21 | 0.14 | 0.125 | -0.44 | 0.14 | 0.002 |
| Age x maternal cond. | -0.15 | 0.15 | 0.303 | -0.16 | 0.15 | 0.265 |
| **Nondominant-sired** |  |  |  |  |  |  |
| Intercept | 14.93 | 0.79 | <.001 | 18.77 | 0.79 | <.001 |
| Litter size | 0.09 | 0.47 | 0.840 | -0.52 | 0.49 | 0.283 |
| Sex ratio (% male) | -0.51 | 0.87 | 0.557 | 1.20 | 0.93 | 0.196 |
| Dom-sired | 2.63 | 1.07 | 0.014 | 3.36 | 1.07 | 0.002 |
| Control-sired | 0.86 | 0.81 | 0.291 | 0.32 | 0.81 | 0.694 |
| Maternal control | 0.80 | 0.89 | 0.371 | 0.06 | 0.89 | 0.944 |
| Age | 2.28 | 0.15 | <0.001 | 3.10 | 0.15 | <.001 |
| Age x litter size | -0.00 | 0.09 | 0.967 | -0.09 | 0.10 | 0.349 |
| Age x sex ratio | -0.03 | 0.17 | 0.870 | 0.24 | 0.19 | 0.203 |
| Age x dom-sired | 0.23 | 0.20 | 0.260 | 0.40 | 0.20 | 0.047 |
| Age x control-sired | 0.02 | 0.16 | 0.911 | -0.03 | 0.16 | 0.836 |
| Age x maternal cond. | -0.14 | 0.18 | 0.438 | -0.30 | 0.18 | 0.088 |
| **Control-sired** |  |  |  |  |  |  |
| Intercept | 15.79 | 0.20 | <.001 | 19.09 | 0.19 | <.001 |
| Litter size | -0.03 | 0.13 | 0.806 | -0.29 | 0.13 | 0.023 |
| Sex ratio (% male) | -0.01 | 0.11 | 0.962 | -0.17 | 0.12 | 0.152 |
| Dom-sired | 1.77 | 0.75 | 0.019 | 3.04 | 0.75 | <.001 |
| Nondom-sired | -0.86 | 0.81 | 0.291 | -0.32 | 0.81 | 0.694 |
| Maternal control | -1.46 | 0.48 | 0.002 | -3.31 | 0.47 | <.001 |
| Age | 2.30 | 0.04 | <.001 | 3.07 | 0.04 | <.001 |
| Age x litter size | 0.04 | 0.03 | 0.124 | -0.23 | 0.04 | <.001 |
| Age x sex ratio | 0.04 | 0.02 | 0.041 | -0.01 | 0.02 | 0.774 |
| Age x dom-sired | 0.21 | 0.14 | 0.125 | 0.44 | 0.14 | 0.002 |
| Age x nondom-sired | -0.02 | 0.16 | 0.911 | 0.03 | 0.16 | 0.836 |
| Age x maternal cond. | -0.23 | 0.09 | 0.012 | -0.68 | 0.09 | <.001 |

Selected outputs from full before-puberty model separated by offspring sex and paternal condition. Paternal factors are shaded in gray. Age centered at six weeks. Litter size centered at median of eight pups. Percent of litter that is male is centered at median of 50%. Parental effects of social experience are in comparison to control-sired offspring. Random effects include animal ID and birthcage. B = coefficient.

**SI Table 4.1.** Paternal status model. Top 15 genes, sorted by posterior probability.

| Gene | description | Bayes factor | Posterior probability | Effect (Dom vs. Non) |
| --- | --- | --- | --- | --- |
| Snora21 | small nucleolar RNA H/ACA box 21 | 197.767 | 0.956473 | -0.000562205 |
| Gm21738 | predicted gene 21738 | 191.871 | 0.955195 | -0.172187 |
| Gm24336 | predicted gene 24336 | 164.244 | 0.94805 | -0.0398351 |
| Gm43042 | predicted gene 43042 | 100.012 | 0.91744 | 0.0846968 |
| Vaultrc5 | vault RNA component 5 | 83.6057 | 0.902814 | -0.478314 |
| Gm45104 | predicted gene 45104 | 82.4589 | 0.901595 | -0.992521 |
| Gm42755 | predicted gene 42755 | 56.1143 | 0.861782 | 0.000300011 |
| Pcbp1 | poly(rC) binding protein 1 | 55.6301 | 0.860746 | -0.998963 |
| Gm11337 | predicted gene 11337 | 46.0112 | 0.836397 | -0.524529 |
| Pcca | propionyl-Coenzyme A carboxylase alpha polypeptide | 43.4249 | 0.828326 | -0.620517 |
| Hnrnpa3 | heterogeneous nuclear ribonucleoprotein A3 | 30.835 | 0.774068 | -0.822176 |
| Pabpn1 | poly(A) binding protein nuclear 1 | 26.0572 | 0.743277 | -1.03724 |
| G6pc | glucose-6-phosphatase catalytic | 23.0107 | 0.718844 | 0.735126 |
| Nop10 | NOP10 ribonucleoprotein | 21.3924 | 0.703873 | 0.498622 |
| Prkar1a | protein kinase cAMP dependent regulatory type I a | 20.7502 | 0.697481 | -0.499644 |

**SI Table 4.2.** Paternal rank model. Top 15 genes, sorted by posterior probability.

| Gene | description | Bayes factor | Posterior probability | Effect size (rank) |
| --- | --- | --- | --- | --- |
| Pcbp1 | poly(rC) binding protein 1 | 55.6301 | 0.860746 | -0.998963 |
| Pcca | propionyl-Coenzyme A carboxylase alpha polypeptide | 43.4249 | 0.828326 | -0.620517 |
| Hnrnpa3 | heterogeneous nuclear ribonucleoprotein A3 | 30.835 | 0.774068 | -0.822176 |
| G6pc | glucose-6-phosphatase catalytic | 23.0107 | 0.718844 | 0.735126 |
| Nop10 | NOP10 ribonucleoprotein | 21.3924 | 0.703873 | 0.498622 |
| Prkar1a | protein kinase cAMP dependent regulatory type I a | 20.7502 | 0.697481 | -0.499644 |
| Ndufs7 | NADH dehydrogenase (ubiquinone) Fe-S protein 7 | 17.1771 | 0.656188 | 0.366851 |
| Hpn | hepsin | 16.475 | 0.646713 | 0.334252 |
| Herpud1 | homocysteine-inducible ER ubiquitin-like domain 1 | 15.392 | 0.631026 | 0.726326 |
| Rpl14 | ribosomal protein L14 | 10.8812 | 0.547311 | 0.267357 |
| Hnrnpd | heterogeneous nuclear ribonucleoprotein D | 10.4815 | 0.538022 | -0.993686 |
| Gm29216 | predicted gene 29216 | 8.00888 | 0.470864 | 0.468597 |
| Arg1 | arginase liver | 7.84042 | 0.465571 | 0.340062 |
| Rn18s-rs5 | 18s RNA, related sequence 5 | 7.42251 | 0.451972 | 0.0402687 |
| Egfl7 | EGF-like domain 7 | 7.09803 | 0.440925 | 0.663961 |

